# Identifying representational structure in CA1 to benchmark theoretical models of cognitive mapping

**DOI:** 10.1101/2023.10.08.561112

**Authors:** J. Quinn Lee, Alexandra T. Keinath, Erica Cianfarano, Mark P. Brandon

## Abstract

Decades of theoretical and empirical work have suggested the hippocampus instantiates some form of a cognitive map. Yet, tests of competing theories have been limited in scope and largely qualitative in nature. Here, we develop a novel framework to benchmark model predictions against observed neuronal population dynamics as animals navigate a series of geometrically distinct environments. In this task space, we show a representational structure in the dynamics of hippocampal remapping that generalizes across brains, discriminates between competing theoretical models, and effectively constrains biologically viable model parameters. With this approach, we find that accurate models capture the correspondence in spatial coding of a changing environment. The present dataset and framework thus serve to empirically evaluate and advance theories of cognitive mapping in the brain.

**HIGHLIGHTS:**

- We identify representational structure in CA1 remapping that is reliable across brains.
- We directly compare models of cognitive mapping to this representation in CA1.
- Models based on local boundary distance and direction predict CA1 representation.
- This approach reveals a biologically viable parameter space for model predictions.
- Accurate models capture the correspondence of spatial codes across environments.

## INTRODUCTION

The hippocampus and associated regions of the neocortex are thought to support diverse learning and memory processes through cognitive maps instantiated in the activity of principal neurons^1^. Growing evidence suggests that such maps are formed across metric spaces to support flexible, goal-directed navigation and long-term memory^2,3^; functions that are impaired following hippocampal damage^4–7^. These findings have recently motivated an increasing number of theories expressed in computational models to explain how the hippocampal system instantiates cognitive maps^8^. However, there has been no consensus on how to compare predictions across models and, importantly, against empirical observation.

To further our understanding of the organization of neural codes throughout the brain, we require experiments to make quantitative comparisons across regions, assays, and theoretical models. One efficient framework toward this end is to collect large datasets in common task spaces where changes in neural representation can be robustly quantified across subjects, and directly compared to model predictions^9^.

A common approach to quantify the determinants of spatial coding is through “remapping”, wherein changes to a familiar environment’s sensory or behavioral conditions change the spatial tuning of selective neural populations. The pattern of remapping observed in changing environments is thought to reflect how the brain disambiguates and relates spatial-contextual information in a changing space, which is likely important for learning and memory processes^10–12^. Despite the widespread assumption that remapping reflects a common representational structure across animals, no studies have examined the reliability of remapping dynamics across subjects at the population level.

While the hippocampus remaps following changes to a wide range of behavioral, sensory, and task conditions, environmental geometry is among the strongest determinants of hippocampal coding^13^. Indeed, extensive behavioral studies have shown that geometry predicts navigation behavior across species^14,15^, and neurophysiological studies further demonstrate that spatial codes across the hippocampal system remap following changes to environment shape^12,13,16–19^. Geometric deformation is a common experimental approach to examine these determinants of neural coding, in which environmental boundaries are added, removed, or shifted. Importantly, geometry can be parametrically varied to create condition-rich experimental designs in which theories of cognitive mapping make competing predictions. Popular theoretical views of cognitive mapping differ in how each posits the hippocampal system constructs a representation of space based on globally allocentric or local environmental features and how cognitive maps might be learned from behavior and experience.

To address this need, we recorded from large populations in hippocampal subregion CA1 in a condition-rich geometric deformation paradigm (5,413 unique neurons across 207 sessions in 10 geometries, forming 69,744 rate maps). Leveraging a representational-similarity framework, we show that remapping in a geometrically dynamic environment reflects a common representational structure that is highly reliable across subjects and find that experience increases the reliability of remapping in CA1. We then demonstrate that these data discriminate between competing theoretical model predictions and can be used to constrain neurobiologically viable model parameters, wherein accurate predictions capture the relationship in spatial coding of a changing environment. The present dataset and approach thus provide the first quantitative benchmark for theoretical advances in cognitive mapping research.

## RESULTS

### Cognitive mapping of a geometrically dynamic environment across protracted experience

Geometric deformation procedures are typically designed to examine changes in neural coding and behavior between two geometries (e.g., square versus circle), requiring the navigator to explore each environment or linear “morphs” between two reference geometries^12,13,20–22^. While this approach has been instrumental to uncover how geometry determines behavior and neural coding, model predictions can be more clearly distinguished by increasing the number and diversity of deformations performed in a single task.

To this end, we partitioned an open square (75 x 75 cm) into a 3 x 3 grid-space to allow systematic manipulations of environmental geometry. We recorded large neural populations in CA1 with miniscope calcium imaging while mice freely explored a sequence of 10 geometrically distinct environments (Figure 1A-B, S1)^23,24^. Importantly, we selected these distinct geometries to induce strong remapping within the same environment across the entire CA1 population. Following habituation to the square arena, animals were exposed to a random sequence of geometries across days that started and ended with the square. The same sequence was repeated up to three times within animals but was randomized across mice (Figure 1C-D).

**Figure 1.**
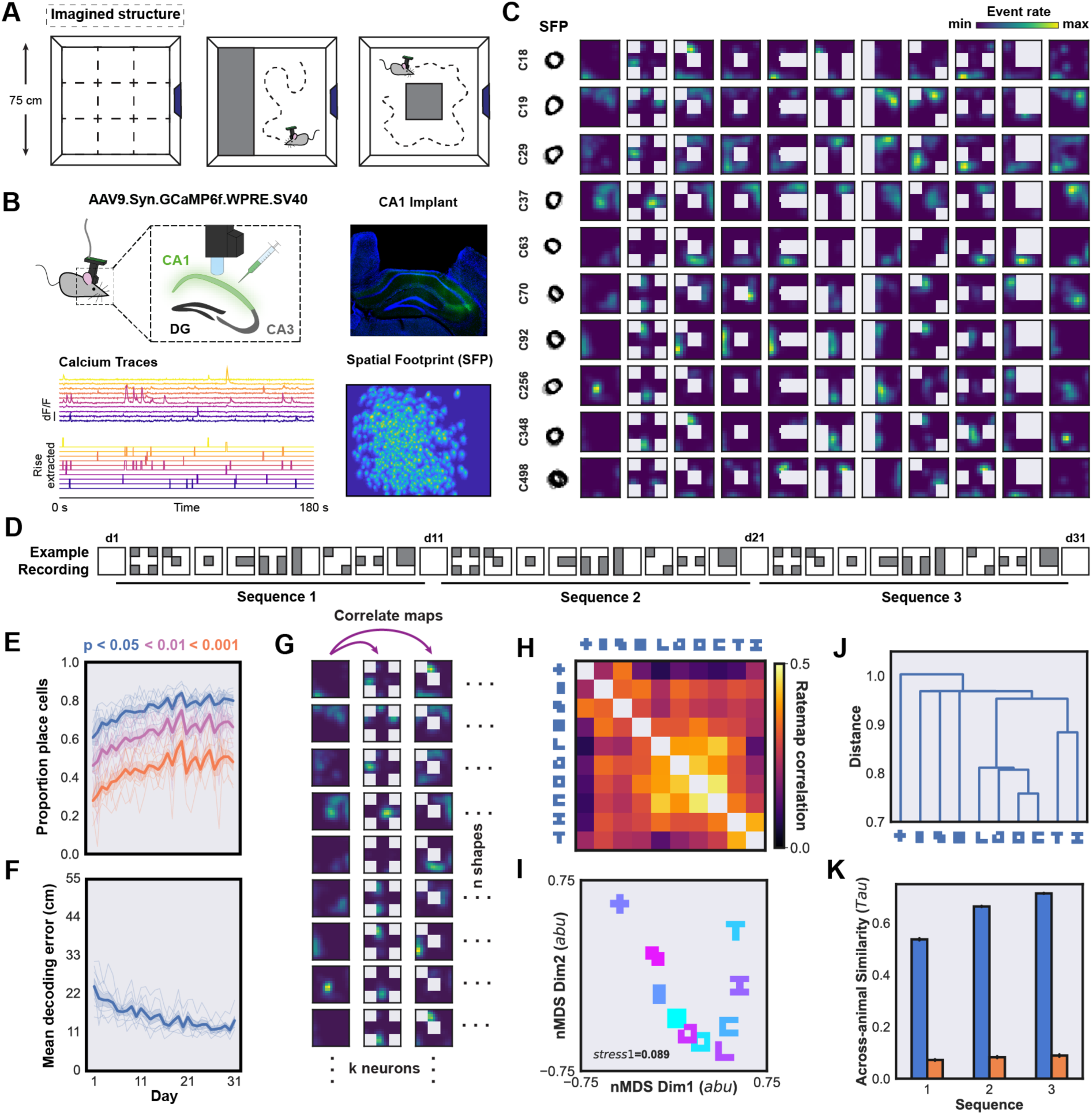
The representational dynamics of remapping in CA1 following geometric deformation is reliable across brains (**A**) To perform systematic manipulations of environmental geometry, we treated a square open field (75 x 75 cm) as an imagined 3 x 3 grid structure, blocking select partitions of the grid with 25 cm walls to create unique geometries in a familiar environment while mice freely navigated the open field. (**B**) To record large neuronal populations in CA1, mice were injected with a viral construct to express GCaMP6f in dorsal CA1 and implanted with a gradient-refractive index (GRIN) lens targeting CA1 (Methods). Spatial footprints and calcium transients were processed and binarized to create an event rate vector for all recorded cells. (**C**) Example rate maps of randomly selected cells registered on all days from a single sequence. The outline of aligned spatial footprints for each cell are shown (left) with rate maps for a target cell across deformations scaled to the maximum of rate maps across days (right). (**D**) Following pre-exposure to the environment, we recorded a repeating sequence of randomly ordered geometries across days, starting and ending with the square environment for each sequence for up to three total repetitions (31 days). (**E**) The proportion of cells that were split-half reliable (*P*<0.01) for each animal and the group average across all recorded sessions. We observed a significant increase in the proportion of cells identified as place cells across sessions (ANOVA: *P*<0.0001; *F*=37.5825). The proportion of place cells according to different p-value thresholds are shown in each color. The group average is shown in the bold trace with shaded SEM, while individual animals are shown in thin traces. (**F**) Average Bayesian position decoding error (cm) from all registered cells and animals across all recorded sessions. We observed a significant decrease in decoding error across recorded sessions (ANOVA: *P*<0.0001; *F*=7.9845). (**G**) To measure the extent of remapping across all pairs of geometries, we performed a rate map correlation for every cell registered across sessions in each pair of environments, and then calculated the average result across all cells and animals. (**H**) The similarity matrix shows the result of the average rate map correlation across all cells, sessions, and animals. (**I**) To visualize the similarity of all shapes captured in the previous comparison (panel H), we embedded the resulting similarity matrix in two-dimensions with nMDS in arbitrary units (abu), which reveals how remapping dynamics cluster based on geometric features of a familiar environment. The inset stress score indicates the quality of fit of the nMDS embedding according to Kruskal stress (*stress1*=0.089). (**J**) The dendrogram similarly shows the clustering of environmental geometries based on their rate map similarity, wherein shapes with a smaller distance are more similar to one another, and those with greater distances differ more greatly in their remapping pattern. (**K**) To measure how similar the pattern of remapping is at the population-level across individual animals, we measured the rank-order correlation of similarity matrices calculated for each animal. The blue bar indicates the true across-brain similarity of remapping, while the orange bar shows the average similarity of shuffled matrices for each sequence. While a two-way ANOVA revealed that the across-brain similarity was significantly above chance (*P*(Shuffle)<0.0001, *F*(Shuffle)=1725.2393), we also found a significant effect of sequence (*P*(Sequence)<0.0001, *F*(Sequence)=37.3110), and sequence x shuffle interaction (*P*(Sequence X Shuffle)<0.0001, *F*(Sequence X Shuffle)=1151.1242) wherein the representational dynamics of remapping becomes more similar across brains with experience.

To measure the quality of spatial coding across recordings, we calculated the split-half spatial reliability for registered cells and position decoding error with a naive Bayesian method^25,26^. Across one month of recording, we observed an increase in the split-half reliability of recorded CA1 populations, increasing the proportion of cells identified as place cells (Figure 1E, S2). In keeping with this result, we also found a significant decrease in position decoding error across recordings, achieving the maximum decoding accuracy reported in recent work (Figure 1F, S2)^27^. These results demonstrate that our CA1 recordings offer a high degree of spatial reliability at the single-cell and population level and motivated the inclusion of all cells in subsequent analyses (mean number of cells per animal = 773 ± 68 SE, minimum cells per animal = 515, maximum cells per animal = 952).

Next, we asked if the pattern of remapping in CA1 during environmental deformations reflected a common representational structure consistent across brains. To do so, we leveraged a Representational Similarity Analysis (RSA) framework. RSA proposes that measurements of a neuronal representation be transformed into a distance matrix that captures the similarity of all pairwise comparisons between conditions in an experiment (Figure 1G-H)^22,28^. Such distance matrices, which define the structure of the representation^29^, can then be quantitatively compared to uncover if and how representational structure differs. Notably, because distance matrices are agnostic to the measurements from which they were derived, they can be compared across subjects, experimental groups, and theoretical models, which has led to its widespread adoption in human fMRI, EEG, and model comparison research^28–30^. Recent advances in high-yield neuronal recording technology have also enabled its application to study large-scale neuronal dynamics with cellular resolution^22^.

To construct an RSM using the rate map correlation as a measure of similarity, we calculated the Pearson correlation between rate maps in each pair of environments for cells registered across sessions in a geometric sequence (Figure 1G). We then computed the average rate map correlation across all registered cells, sequences, and animals to produce a representational similarity matrix (RSM) of the pairwise similarity of all geometries (Figure 1H). This analysis revealed a wide range of similarities in the spatial mapping across geometries and that the most significant amount of remapping was observed following the omission of corners and walls (Figure 1H, S3). Next, we transformed this RSM into a set of points (one for each environment) in a two-dimensional space where proximity reflects the similarity between each geometry’s rate maps using a technique called non-metric Multidimensional Scaling (nMDS), and further visualized clustering of rate map similarities in a dendrogram (Figure 1J). These analyses show that the similarity of CA1 maps cluster based on shared geometric features of environments.

We then asked to what extent the pattern of remapping in CA1 was reliable across subjects in our task. To do so, we calculated the rank-order correlation of RSMs for each pair of subjects in each sequence and for random permutations of the same RSM (Figure 1K). We observed that the pattern of remapping in CA1 was highly reliable across subjects and that this across-subject reliability grew across sequences, providing the first demonstration to our knowledge that remapping reflects a common representational structure at the population level in CA1.

### A rich and reliable representation of geometric remapping in CA1

While leveraging the classical rate map correlation with an RSA-based framework revealed that the representational dynamics of remapping are reliable across brains, this approach also limits comparisons to allocentrically overlapping areas of environments. Based on our grid-based approach to environmental deformation, the rate map correlation can be more powerfully applied to individual partitions to allow specific and condition-rich comparisons of cognitive mapping within and across environments. Such comparisons can be used to characterize the representational structure of CA1 remapping in a geometrically dynamic environment, to further evaluate the reliability in remapping across brains, and specifically reveal where competing models succeed or fail to predict spatial representation.

To this end, we partitioned each cell’s rate map according to the 3 x 3 grid design in each geometric sequence (25 x 25 cm per partition). For each cell, we then calculated the Pearson correlation coefficient of rate map partitions within and across all geometries (Figure 2A)^22^. This yielded a large (75^2^ including sampled partitions) and expressive representational similarity matrix (RSM) for comparison across repeated experiences within the same animal, across animals, and against model predictions (Figure 2B)^29^.

**Figure 2.**
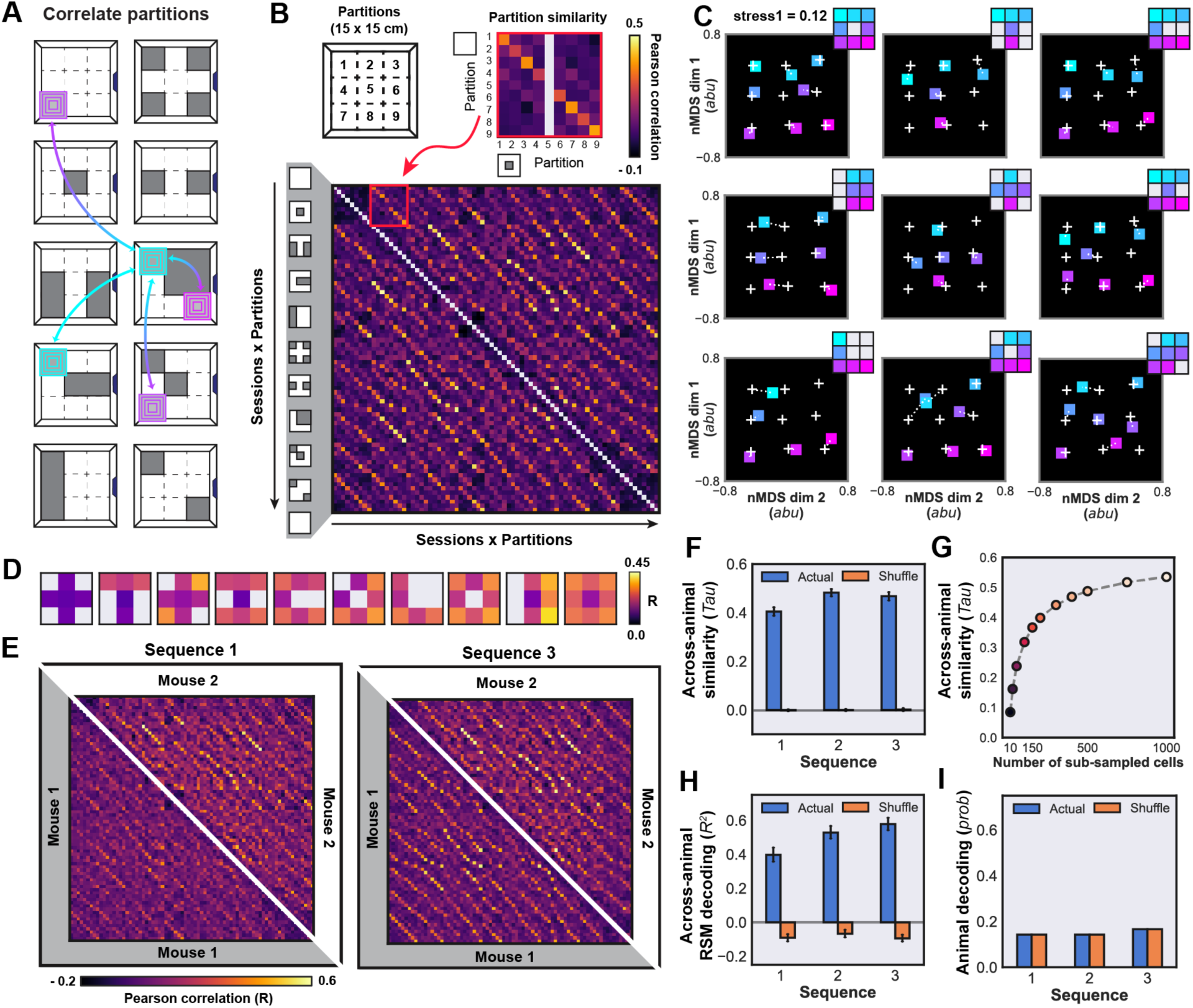
A condition-rich approach to examine the representational structure of remapping with environmental geometry (**A**) The schematic illustrates how partitions within and across geometries are compared in each sequence. (**B**) To construct an RSM for each animal and sequence, we calculated the average Pearson correlation across all pairwise comparisons of partitioned rate maps from matched cells across registered sessions. For each geometry this comparison yields a 9 x 9 matrix of similarities (75^2^ comparisons within a sequence for all sampled partitions). (**C**) The nMDS embeddings of the average RSM reveals the effect of geometric deformation on representational similarity among partitions (*stress1* = 0.12). The average result of the square environment at the start and end of each sequence is shown in white cross points, while dashed white lines show translation in the embedding space of the matched partition relative to the square. Colors indicate true allocentric position of each partition across environments (inset shown at the top right of each panel). (**D**) Average similarity of individual partitions relative to the matched partition in the square environment across all sequences and animals. (**E**) The RSMs from two separate mice are shown in upper and lower diagonal across sequences, wherein the white diagonal line separates the triangle of the RSM for each mouse. (**F**) Rank-order correlation (Kendall’s Tau; mean ± SEM) for all across-animal RSM comparisons within each sequence, compared to rank-order correlation of shuffled RSMs. We observed a significant effect of sequence on animal-wise representational similarity compared to sequence-shuffled controls (ANOVA: *P*(Sequence)<0.0001, *F*(Sequence)=22.3110; *P*(Shuffle)<0.0001, *F*(Shuffle)=7673.5486; *P*(Sequence X Shuffle)<0.0001, *F*(Sequence X Shuffle)=24.7201). (**G**) To measure the relationship between the number of observed cells across sessions and the similarity in representational structure across animals, we sub-sampled cells 1000 times and re-computed the average similarity across animals. This analysis revealed that the high level of similarity can be observed only with hundreds of neurons recorded across sessions. (**K**) Decoding of RSMs across animals for all partitions and geometries (ANOVA: *P*(Sequence)<0.0001, *F*(Sequence)=16.3418; *P*(Shuffle)<0.0001, *F*(Shuffle)=1651.4348; *P*(Sequence X Shuffle)<0.001, *F*(Sequence X Shuffle)=11.7739). (**I**) Decoding probability of animal identity from the respective RSM across sequences compared to a shuffled control. We did not observe a significant effect of shuffling animal identity on RSM decoding (Chi-square: *P*=1.000, *χ*2=0.000).

To visualize the differences in hippocampal representational structure across geometries, we first averaged our RSMs across all sequences and animals (Figure 2B). Next, we transformed this averaged RSM into a set of points (one for each partition and geometry) in a two-dimensional space with nMDS^31^, which allows us to visualize a high-dimensional representation (RSM with 75^2^ dimensions) in a digestible lower-dimensional space while preserving the relative similarity between points. This embedding revealed that occluding one partition resulted in nearby partitions becoming more similar to the occluded area (Figure 2C). To quantify the effect of deformation in each partition, we measured the correlation of each partition relative to the square environment (Figure 2D). This revealed a strong effect of partition on similarity, wherein corners of the environment exhibited the greatest similarity, and the center partition showed the least similarity across geometries (Figure 2D, S4A). Interestingly, this pattern mirrors the effect of corner omission on overall map similarity in previous analyses, wherein corner omission resulted in the greatest remapping (Fig 1I, S3). Comparing all geometries to the first square of each sequence, we also found a strong effect of geometry on overall similarity, in which the cross– and T-shaped geometries showed the least similarity (i.e., the strongest deformation), while the rectangle and subsequent square geometries showed the greatest similarity (Figure S4B). These results demonstrate that our population-level RSM effectively captures the impact of geometric deformations on hippocampal representational structure.

To compare the reliability of this representational structure within and across brains, we computed the average RSM across cells within each animal for each sequence. Comparison of these RSMs revealed a high degree of similarity in representational structure across animals (Figure 2E-F). We also found that the similarity of representations across animals significantly increased with experience (Figure 2E-F). Given the large number of cells recorded with our approach, we also examined how the reliability of representation across animals was related to the number of cells recorded across a geometric sequence. To do so, we randomly sub-sampled cells from the recorded population, and re-computed the average similarity across subjects. This analysis revealed that the reliability of remapping across subjects we observed requires hundreds of cells to be recorded across experimental conditions (Figure 2G), underscoring the unique advantages of combining longitudinal population recording with condition-rich task design.

Next, we asked if we could decode animal identity from the RSM based on reliable individual differences, and whether the RSM could be accurately predicted across animals and sequences. We anticipated that if the RSM is highly reliable across brains and individual differences are noise-driven rather than reliably subject-specific, then we should not be able to predict animal identity from its RSM. By contrast, if dissimilarities across subject’s RSM are driven by reliable individual differences, then animal identity should be decodable from the RSM. While the RSMs can be accurately predicted across animals and sequences, we were not able to decode animal identity (Figure 2H-I). Further analysis of male and female subjects also confirmed there are no sex differences in the CA1 neural representation in our task (Figure S5). These findings demonstrate that the geometric determinants of the CA1 representation are highly reliable within and across brains.

### Predicting the representational structure of remapping with heuristic models of spatial representation and behavior

A classical view of cognitive mapping posits that the representation of space in the hippocampus is globally-allocentric, and spatial coding varies with Euclidean proximity^1^. By contrast, subsequent observations of remapping suggested that the hippocampus instead represents locations based on the allocentric relationships between local environmental features, such as walls or boundaries^13^. Recently, a growing body of work has also suggested that the hippocampus forms predictive maps learned through experience and the structure of behavior (e.g., spatial navigation)^32^. We thus sought to address if our condition-rich RSA framework could resolve how well heuristic models of these concepts predict the pattern of remapping we observe in CA1.

We thus constructed RSMs for our task space according to similarity metrics that express the core predictions of each heuristic model. To generate globally-allocentric predictions, we measured the Euclidean distance between all partitions across a geometric sequence, and then transformed each distance to a similarity measure by subtracting from one after normalizing to the maximum distance across environments (Figure 3A). To calculate the similarity of local allocentric boundaries in each partition across geometries, we first described each partition with a one-hot vector where a value of 1 indicates the presence of a boundary, and zero the absence of a boundary for the cardinal directions surrounding a partition (Figure 3B). To calculate the similarity of boundary conditions across partitions, we calculated the hamming distance between one-hot vectors describing each partition’s local boundary condition and subtracted from the maximum. Finally, to measure the similarity of animals’ navigation trajectory in each partition, we first computed a transition matrix for an animal’s position in all locations (5 cm^2^ spatial bins @ 2 Hz; Figure 3C). To measure trajectoral similarity across partitions, we then calculated the rank-order correlation (Kendall’s Tau) between transition matrices across all partitions.

**Figure 3.**
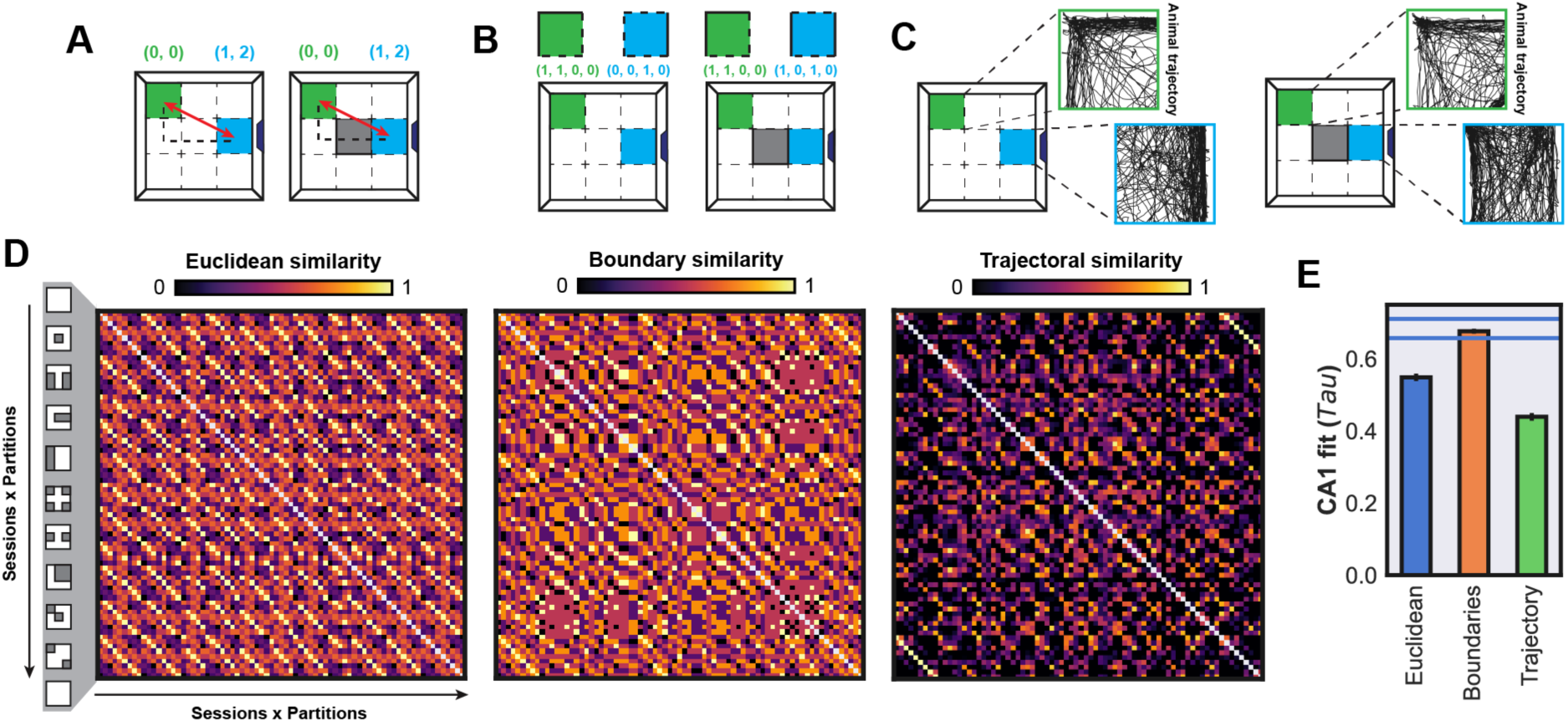
Predicting representational structure with heuristic models of cognitive mapping (**A**) To capture representational structure in our task according to a classical globally-allocentric view of cognitive mapping, we calculate the Euclidean distance between each pair of partitions across all geometries and measured their similarity as the normalized difference between the maximum and actual difference in Euclidean space. (**B**) An alternative view of cognitive mapping is that spatial codes are constructed from on the allocentric location of local boundaries in an environment. To express this view in a heuristic model we constructed a one-hot vector for each partition indicating the location of west, north, east, and south boundaries around each partition, respectively, and computed the hamming distance between the boundary one-hot vectors. To convert this distance to similarity, we then subtracted the distance from the maximum possible distance. (**C**) An emerging view of cognitive mapping is that the structure of cognitive maps is learned from predictive relationships between locations or stimuli in environments that are experienced from the structure of behavior. To illustrate this view in our task, we measured structure of navigation trajectories by constructing a transition matrix between spatial locations (5 cm^2^ spatial bins @ 2 Hz) for each partition. We then calculated the rank-order correlation (Kendall’s Tau) between transition matrices across all partitions and environments. (**D**) Using each heuristic model and measure of similarity, we constructed RSMs to predict the representational structure of cognitive mapping across all environments and partitions, as we performed with our CA1 data. The RSMs illustrate differences in the pattern of remapping that emerges from each heuristic model of cognitive mapping in our task. (**E**) To examine how well each model predicts the pattern of remapping in CA1, we measured the rank-order correlation (Kendall’s Tau) between each predicted RSM and CA1, which revealed that while each account predicts a significant amount of variation in representational dynamics of CA1 in our task (*P*<0.0001), a one-way ANOVA revealed a significant effect of model on prediction accuracy (*P*(Model)<0.0001, *F*(Model)=13981.1166). Namely, the heuristic boundary-based prediction offers the maximum level of explanation within the calculated noise ceiling (STAR Methods).

This analysis revealed that each heuristic model generates distinct predictions in our task. For example, the partitions of greatest similarity in the classic globally-allocentric model predictions (same Euclidean location), were often not the regions of greatest similarity in the allocentric boundary or trajectory-based model predictions (Figure 1D). To measure how well each model predicts CA1 representation in our task, we measured the rank-order correlation between each model RSM against that observed in CA1. We found that, while each heuristic model alone predicted a significant amount the CA1 representation, there was a significant difference between heuristic models and the similarity of local allocentric boundary locations best predicted CA1 representation (Figure 3E; STAR Methods). Given these outcomes, we then sought to address if neurobiological models that instantiate each view would similarly predict the pattern of remapping in CA1.

### Benchmarking neurobiological models of cognitive mapping with neuronal population data

Our experimental and analytic approach was designed to provide a highly reliable measure of the representational structure of hippocampal cognitive mapping and remapping. Having addressed this need, we explored the possibility that these data can be used to benchmark heuristic model predictions of cognitive mapping, which revealed that local allocentric boundary conditions best predicted the pattern of remapping observed in CA1. We then asked if neurobiological models that express globally-allocentric, allocentric boundary locations, or behaviorally driven dynamics would similarly predict the pattern of remapping we observe in CA1. Specifically, we sought: (1) quantification of neurobiological model prediction accuracy against an observed neuronal representation in the brain, (2) discrimination between model predictions in a common task space, and (3) empirical constraints for biologically viable model parameters.

While comparison among all models of cognitive mapping is beyond the present scope, we chose to assay several popular neurobiological models of cognitive mapping that instantiate the core predictions of the heuristic models described above. To capture the globally-allocentric hypothesis of cognitive mapping, we generated predictions from a naïve model of globally allocentric place fields (PC), in addition to the popular grid-to-place model (GC2PC)^33^. To contrast competing predictions on remapping dynamics following geometric deformation, we implemented a boundary-tethered grid-to-place model (bt-GC2PC)^17^ and a boundary-vector-to-place model (BVC2PC)^34–36^. While the bt-GC2PC model predicts remapping in CA1 spatial coding due to translations of grid cell receptive fields upon boundary contact, the BVC2PC model instead predicts remapping is driven by a specific boundary-vectoral representation of environment geometry. Finally, a growing number of theoretical proposals suggest that the hippocampus represents a predictive map that captures the temporal structure of behavior (e.g., navigation) in the dynamics of learning and remapping^32^. Importantly, the description of behavioral states in such models can be defined in any frame of reference, such as a globally-allocentric or boundary-vector space^37,38^. To express behaviorally-driven predictions on cognitive mapping and remapping in CA1, we implemented a globally-allocentric place cell to successor feature model (PC2SF)^32,37^ and a boundary-vector to successor feature model (BVC2SF)^32,37,38^.

According to each view, we simulated predicted neural dynamics in a population of model neurons as an artificial agent navigated our paradigm following the actual trajectories for each animal (Figure 4A; STAR Methods)^39^. These model neurons formed a basis set from which hippocampal model predictions were derived (Figure 4A). Treating these model neurons as we had treated recorded populations in CA1, we computed model RSMs for each geometric sequence and animal (Figure 4B-C, S5). *Prime facie*, model RSMs varied considerably in this task space, suggesting this approach provides an expressive characterization with which to adjudicate between predictions. We then measured the rank-order correlation of each model RSM against the recorded CA1 representation, which revealed significant differences in the ability of each model to account for the CA1 representational structure we observed (Figure 4D). This approach circumvents challenges to determine correspondence between empirically observed and simulated neural activity^28^. We found that the BVC2PC model provided the greatest level of prediction accuracy, achieving the theoretical maximum level of prediction (noise ceiling; STAR Methods) (Figure 4D). Indeed, the performance of such neurobiological models of cognitive mapping agrees with the outcomes of our previous heuristic model comparisons (Figure 3). Importantly, we also found that successor features learned from globally-allocentric, or boundary-vector features afforded a similar relative level of explanation as comparable models where features are not learned, such as the PC or GC2PC and BVC2PC models. These outcomes suggest that biologically plausible learning rules and behavior can be used to learn features that resemble CA1 representation from a specific (BVC) basis set. However, the addition of behavior alone does not add significantly to a description of cognitive mapping in CA1, and instead the description of agent state (PC or BVC basis) is a greater determinant of model prediction accuracy.

**Figure 4.**
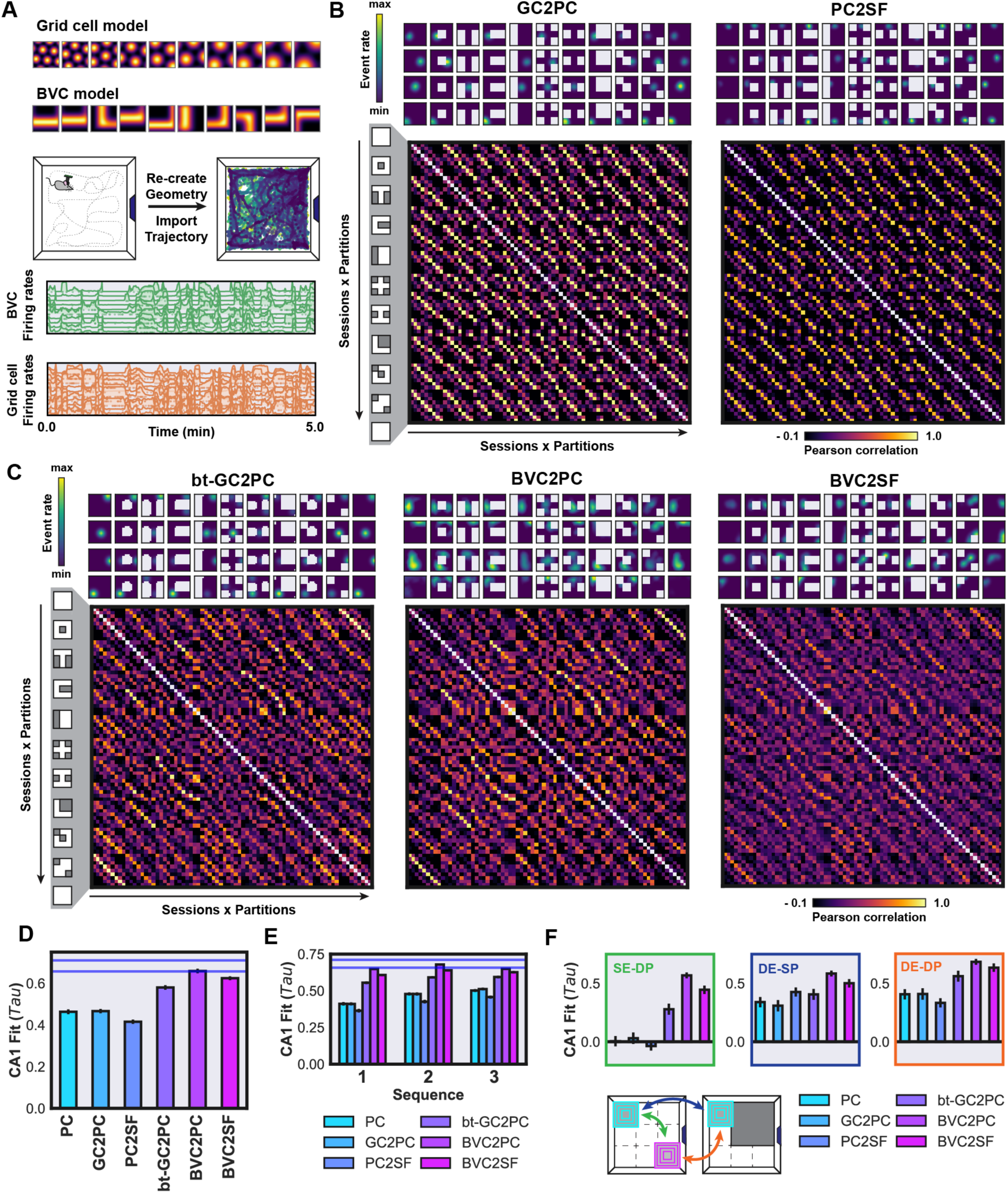
Benchmarking neurobiological model predictions against representational structure in CA1 (**A**) Illustration of firing rate simulation for model features in the parahippocampal system, wherein receptive fields are drawn according to each model, and firing rates are simulated with recorded animal trajectories using an open-source toolbox^39^. Rate maps were constructed from model firing rates for all features and used for model RSM calculation identical to CA1 data. (**B**) Example rate maps and respective RSMs are shown for the globally allocentric GC2PC model, and the globally-allocentric, behaviorally driven PC2SF model. Rate maps show a high level of stability in the GC2PC model across all environments, and behaviorally driven plasticity with a temporal-difference learning rule in the PC2SF model. The bold yellow reveals a high level of similarity for matching partitions in a globally-allocentric reference from across environments in both models. (**C**) Example rate maps and respective RSMs illustrate remapping dynamics across environments for the bt-GC2PC, BVC2PC, and behaviorally driven BVC2SF model. Each panel illustrates the marked difference in remapping patterns compared to the globally-allocentric models, with further distinctions based on the type of boundary-driven remapping (bt-GC2PC vs BVC2PC) and behaviorally driven plasticity (BVC2PC vs BVC2SF). (**D**) To evaluate model predictions against the observed representation in CA1, we measured the rank-order correlation between each model RSM and CA1 (Kendall’s Tau). Each bar shows the fit of each model to CA1 data, while error bars show the bootstrap standard deviation of each fit. Blue lines indicate the upper and lower bounds of noise ceiling, which is considered the margin of maximum prediction accuracy given noise in the CA1 data (Methods). We observed a significant effect of model on predicting CA1 representation, with BVC2PC and BVC2SF models showing prediction accuracy within noise ceiling (ANOVA: *P*(Model)<0.0001, *F*(Model)=419.7742). All model predictions predict a significant amount of the representational structure observed in CA1 (*P*<0.0001). (**E**) To examine whether the representational structure of remapping in CA1 changes according to different model predictions with experience, we examined the prediction accuracy of each model against CA1 data for each sequence. We observed a significant effect of model (ANOVA: *P*(Model)<0.0001, *F*(Model)=825.0067), sequence (ANOVA: *P*(Sequence)<0.0001, *F*(Sequence)=5018.9717, and model x sequence interaction (ANOVA: *P*(Model X Sequence)<0.0001, *F*(Model X Sequence)=2047.5743). While the prediction accuracy of models improved across sequences, we observe an increase in the ability of allocentric models to predict CA1 representation in later compared to earlier geometric sequences. However, the same rank-order of model performance is preserved across all sequences, suggesting a convergence in CA1 representation, rather than a change in the pattern of remapping. (**F**) To examine specific patterns of remapping that each model explains, we broke down all comparisons of partitions within and across geometries into three types: same environment – different partition (SE-DP); different environment – same partition (DE-SP); different environment – different partition (DE-DP). A two-way ANOVA revealed a significant effect of model, and comparison type, and model x comparison interaction on prediction CA1 representation (ANOVA: *P*(Model)<0.0001, *F*(Model)=8014.9025; *P*(Comparison)<0.0001, *F*(Comparison)=13718.1338; *P*(Model X Comparison)<0.0001, *F*(Model X Comparison)=895.7919).

Given our previous observation that CA1 representation became more reliable across animals with experience (Figure 1K, 2F), we sought to address whether neurobiological models differently predict CA1 representation in each geometric sequence of our task. Specifically, we aimed to address whether the prediction accuracy of models changed across sequences. To do so, we examined the accuracy of model predictions for each sequence (Figure 4E), which revealed a significant effect of sequence and model. While geometrically driven models (bt-GC2PC, BVC2PC, BVC2SF) afforded a similar level of explanation across sequences, there was an increase in the prediction accuracy of globally allocentric models with experience (PC, GC2PC, PC2SF), which could be driven by within-session increases in spatial reliability that we observed previously (Figure 1E, 1F). Importantly, the relative order of model performance did not change across sequences in our experiment, suggesting a convergence in spatial representation with experience for which boundary-vector based models afforded accurate predictions.

We next asked whether the success of each model to predict the structure of CA1 representation was due to a subset of predictions in our task. Specifically, we asked how well each model predicted remapping across different partitions within a single environment (SE-DP; Figure 4F), the same partition in different environments (DE-SP; Figure 4F), or different partitions of different environments (DE-DP; Figure 4F). To this end, we calculated the rank-order correlation of each model with CA1 data for each subset of comparisons. This analysis revealed that BVC-based models best accounted for the pattern of remapping observed across partitions within environments (SE-DP), the same partitions in different environments (DE-SP), and different partitions in different environments (DE-DP). Notably, we found that globally-allocentric models (PC, GC2PC, PC2SF) failed to explain the remapping across partitions within an environment (Figure 4F), confirming that spatial coding within complex geometries is non-Euclidean, and likely based on local environmental features and geometric motifs. Indeed, we notice repetitions of place fields in geometrically similar regions within environments (Figure 1C). We also note that boundary-driven and globally-allocentric models make more similar predictions in the comparison of the same partitions across different environments (DE-SP), suggesting that predictions in these spaces are less distinct, while the comparison of different partitions within and across geometries better distinguish model predictions (SE-DP, DE-DP).

Given that the BVC2PC model provided the highest quality fit to the CA1 representation in one parameterization, we next asked whether model fit could be used to constrain a biologically plausible parameter space. To this end, we performed a large sweep of the parameter space for the boundary vector features forming the basis set in the BVC2PC model, including the parameters determining distance-based tuning and the shape of the distribution from which tunings were drawn (STAR Methods). Specifically, we examined the effect of BVC tuning distance constant and scalar (Figure 5A), which both affect width of BVC tuning at their preferred distance at target boundary angles. To manipulate the distribution of preferred BVC tuning distances, we randomly drew preferred tuning distances for each BVC from a beta distribution, controlled by alpha and beta parameters that affect the skewness of the BVC tuning near boundaries (Figure 5B). When beta and alpha values are equal, the preferred tuning distances of BVCs are uniform.

**Figure 5.**
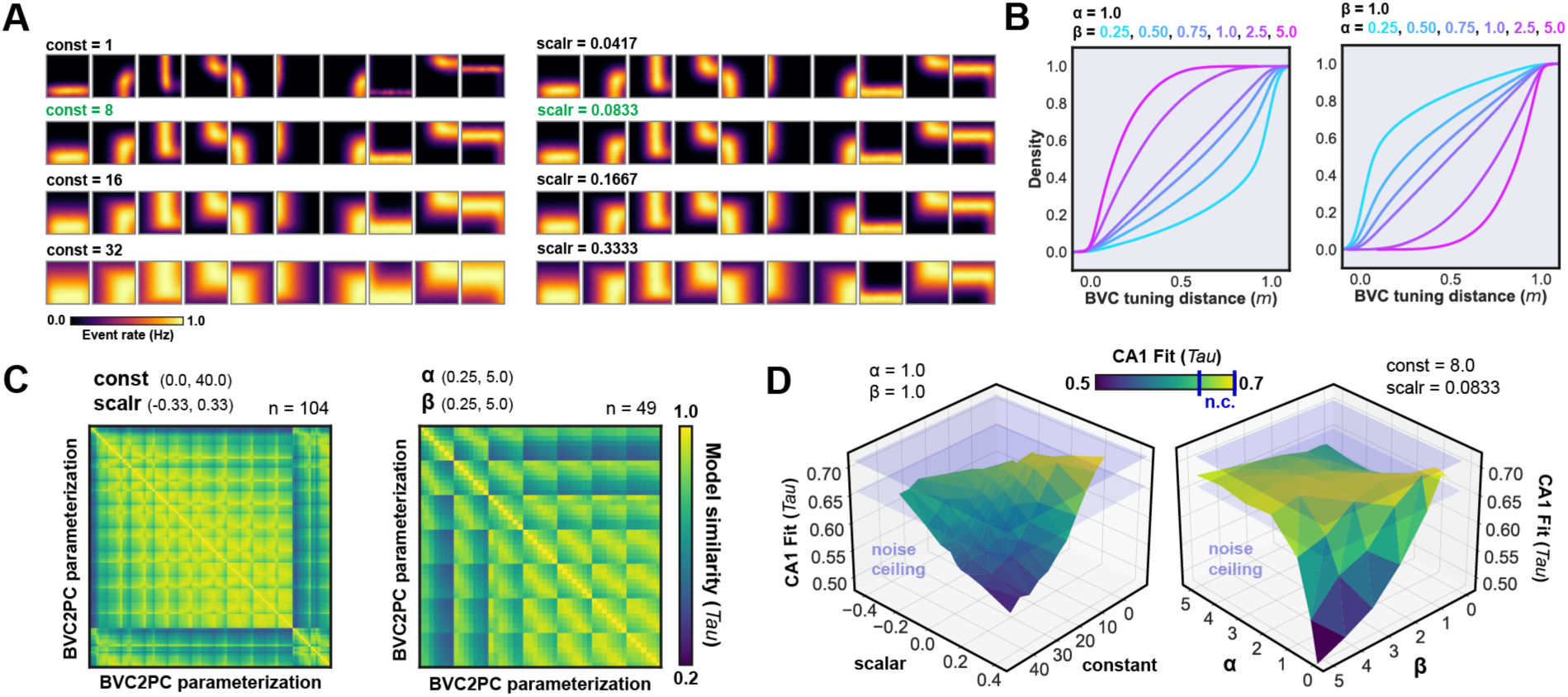
A specific range of model parameters accurately predict CA1 representational structure (**A**) Rate maps illustrate the effect of manipulating the BVC distance constant (left) and distance scalar (right) values in the BVC basis set used for the BVC2PC and BVC2SF model, which differently affect the tuning specificity of BVC model features with distance from boundaries at a specific angle. A model BVC with a specific set of angular tuning parameters is shown in each column, and the manipulated parameters are shown throughout each row. (**B**) The cumulative density plots show the distribution of preferred BVC tuning distances when manipulating the beta (left) and alpha (right) parameters of a beta-distribution. When the two values are equal, this results in a uniform distribution of tuning parameters, while unequal values differently affect skewness of the distribution to have preferred tuning distances near or far from environment boundaries. (**C**) The model-wise RSM shows the measured similarity (Kendall’s Tau) of grid-search parameterizations of the BVC distance constant and scalar (left; *N*=104), and BVC alpha and beta parameters that determined the BVC distance distribution (right; *N*=49). (**D**) The surface plots illustrate the CA1 prediction accuracy landscape of the BVC2PC model with the same parameters shown in (**C**), both for the BVC distance constant and scalar (left), as well as the BVC preferred distance distribution (right). The figures illustrate that a specific range of BVC2PC parameters enters the upper and lower bounds of noise ceiling, revealing a constrained, biologically viable parameter space.

Comparison of model RSMs with each combination of parameters to one another revealed that different parameterizations led to disparate predictions (Figure 5C), demonstrating that our task and RSA approach is sensitive to model parameterization. We then compared all model parameters against CA1 data and found that only a subset of distance tuning and distribution parameters approached the theoretical maximum level of prediction accuracy (Figure 5D, S7). Namely, BVC tuning distances that sampled from an approximately uniform distribution were reliably optimal, with a distance constant of 8 cm and distance scalar value of 12^-1^. These results show the effectiveness of this approach in constraining neurobiologically viable model parameterizations, and in turn allow us to constrain our models using these data prior to application in future experiments.

### Accurate models capture the spatial correspondence of remapping in CA1

A central hypothesis on the role remapping in cognition is to relate spatial-contextual information through the transformation of a neural code. That is, the cognitive map for a location in environment A will correspond to another location in environment A’ following a change to the environment (e.g., geometry) through a specific rule of transformation^10–12^. We reasoned that a precise model of cognitive mapping should therefore capture the spatial correspondence of remapping in a dynamic environment. For example, the BVC2PC model should predict which locations are most and least similar between environment A and A’ when compared to CA1. While the partition-wise RSM comparison coarsely captures this notion, we sought to leverage the large populations recorded in the present study to pursue a detailed comparison of remapping based on this intuition.

To this end, we measured the population vector (PV) correlation in CA1 across all pairs of spatial bins for each pair of geometries, averaged across sequences and animals (Figure 6A). This produced a rich matrix of PV correlations that describe the similarity of all spatial bins to one another within and across environments (Figure 6B). To visualize the correspondence of similar locations across geometries, we selected a seed location in the square geometry to generate similarity maps of all locations in another geometry (Figure 6C). We generated the same PV correlation matrices for model predictions and compared the correspondence of spatial codes in CA1 with each model.

**Figure 6.**
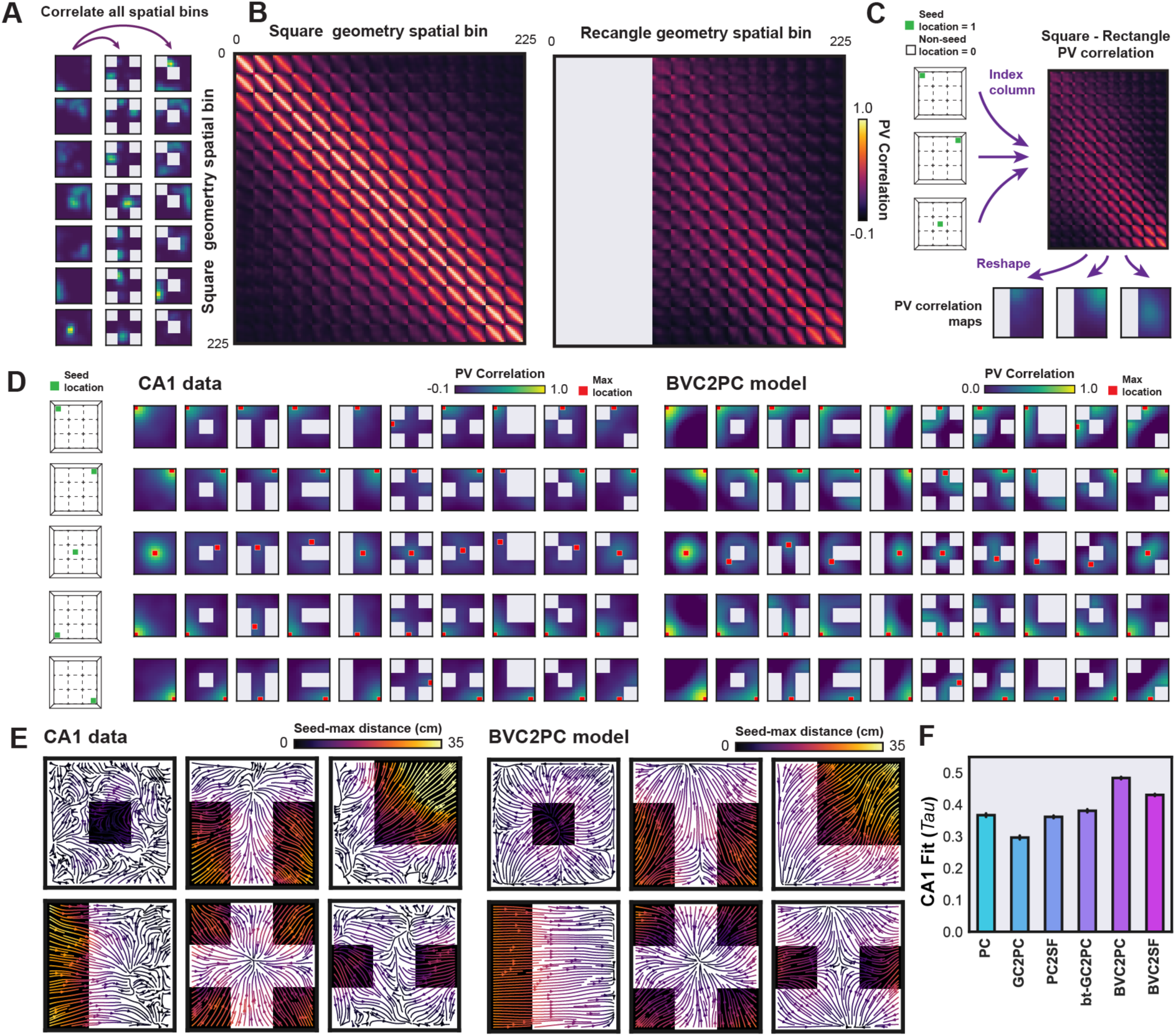
Accurate model predictions capture the spatial correspondence in CA1 remapping across geometries (**A**) To examine the similarity of all locations across geometries following remapping in CA1, we calculated the correlation across all spatial bins in each pair of geometries for every registered, spatially reliable cell. (**B**) The panel shows examples of the correlation structure for two environment comparisons, the correlation of spatial bins in the square environment with each other (left), and the correlation of all square spatial bins with the rectangle geometry (right). (**C**) To reveal the similarity of a single location in the square environment to all locations in the rectangle geometry, we used a single seed location to index and reshape a single column of the pixel-wise correlation matrix. This produces a correlation map of the rectangle environment that shows the similarity of all locations in environment A’ (e.g., rectangle) to a single, seed location of environment A. (**D**) Examples on the left show selected seed locations (corners and center) used to generate spatial correspondence maps organized by rows (right). The spatial correspondence maps show the similarity of all locations to the seed location in the background color, while the location of maximum similarity to the seed is indicated with a red point. CA1 correspondence maps are shown on the left, while those for the BVC2PC model are shown on the right. (**E**) Population vector flow maps show the direction and magnitude of displacement of the maximally correlated population vector in example deformed environments relative to the same location in the square environment. The vector fields reveal the dynamics of remapping across all recorded neurons in the example environments compared to the square environment for CA1 data (left) and the BVC2PC model (right). (**F**) To quantify the similarity of spatial correspondence in remapping of each model with CA1, we measured the rank-order correlation (Kendall’s Tau) between each pair of environments to CA1 data, which revealed a significant effect of model to predict the similarity of population vectors across environments (ANOVA: *P*(Model)<0.0001, *F*(Model)=184.3735).

The visualization of spatial correspondence of select seed locations revealed that the displacement of similar locations in a target environment scaled with the overall level of similarity. We observed a similar pattern in the correspondence of remapped locations in the BVC2PC model with CA1 (Figure 6D). When we visualized the displacement of maximally similar locations to the square environment with population vector-flow maps, we observe a similar pattern of remapping between the BVC2PC model and CA1 data (Figure 6E, S8-9). Direct comparison of the PV correlation matrices of each model with CA1 confirmed that the boundary-vector based models best explained the pattern of spatial correspondence in CA1 remapping across geometries in our task (Figure 6F).

## DISCUSSION

Here we present the first direct quantitative comparison of population level neuronal representation in freely behaving animals against theoretical models of cognitive mapping. Leveraging a similarity-based framework, we show that the representational dynamics of remapping with geometry is highly reliable across brains and increases with experience. We then evaluate the ability of competing theoretical views to predict the pattern of remapping in CA1 and directly compare models of globally-allocentric, local allocentric boundary conditions, and behavioral accounts of cognitive mapping. While each view significantly predicts the pattern of remapping at the population-level in CA1, we find that models based on the allocentric distances and direction of local boundaries best explain the pattern of remapping in CA1. Although behavioral models of cognitive mapping did not result in increased prediction accuracy, such models suggest that hippocampal representations could be learned from a specific basis set and learning rules. Namely, we observed that a constrained range of allocentric, BVC-based models accurately predict CA1 spatial representation (within maximum theoretical limits). The success of such models to predict CA1 representation suggest that an allocentric vector-based code well-describes population level neuronal dynamics in CA1^13,40,41^. Our analysis further reveals that accurate models of cognitive mapping in CA1 capture the correspondence of spatial codes in remapping across changing environments. These are to our knowledge the first results demonstrating that large-scale predictions from neurobiological models can be directly evaluated against population dynamics in freely behaving animals. The present dataset and framework provide a benchmark for theoretical innovations in cognitive mapping research and, more generally, establish a novel approach to compare representational structure across brain regions, assays, species, and theoretical models.

By comparing model predictions in this paradigm, we were able to accurately discriminate between population-level representations with distinct (across-model) and similar bases (within-model parameterizations), suggesting the same approach can be applied to determine the relationship between neuronal codes across brain regions. For example, the representation we observed in CA1 could be quantitatively compared to populations in the entorhinal, subicular, retrosplenial, and further association cortices following a similar approach^42^. This framework holds the promise to allow us to quantitatively derive the transformation of representational structure across brain-wide networks. When combined with causal circuit manipulations, these experiments can further implicate particular mechanisms thought to be selectively responsible for instantiating specific aspects of this representational structure.

While the hippocampal system is known to remap with changes to a wide range of sensory, behavioral, and task conditions, we focused in the present work on the geometric determinants of cognitive mapping. The reason for our approach was two-fold: 1) geometry is known to be among the strongest determinants of hippocampal remapping; 2) distinct theoretical models offer competing predictions to evaluate against empirically observed representational dynamics in CA1. While this work is the first to show that remapping reflects a reliable representational structure across subjects, future work should determine if the same is true for other effectors of remapping across brain regions. One should expect that if an environmental feature is encoded in the activity of a target neural population, then changes to that feature should produce reliable representational dynamics. Building on the present study, future work should systematically manipulate a range of sensory features of environments to drive remapping.

A growing emphasis of recent work in systems neuroscience is to determine how behavioral states and task objectives shape neuronal representation. While the hippocampus has been long appreciated for its role in spatial coding, navigation and memory, an increasing number of studies report hippocampal coding of non-spatial, behaviorally relevant task features, such as rewards, tones, and evidence accumulation^2,3,43,44^. We suggest that future studies build on this approach to compare neuronal codes across regions and against model predictions under distinct behavioral objectives.

In the present work we found that a specific boundary-vector based neuronal code accurately predicted representational structure CA1. However, the models that performed well in the present task may be agnostic to other sensory or behavioral conditions. Through the development of additional tasks and datasets to measure representational dynamics in the hippocampal system, inspired by the design of the present study, future work should evaluate generalized models of cognitive mapping across a range of benchmarks. Such developments upon the present dataset and approach will offer a powerful testbed for theoretical models within and beyond the hippocampus.

## ACKNOWLEDGEMENTS

We would like to thank Tianmeng Xu, Raphael Lavoie, and Kay Cui for their technical assistance with this project. We are grateful to R. R. Rozeske, M. Oulé, M. H. Yaghoubi, T. George, and members of the Brandon lab for their helpful feedback on early versions of this manuscript. This research was funded by the Canadian Institutes of Health Research (project grants #367017 and #377074 awarded to M.P.B.) and the Natural Sciences and Engineering Research Council of Canada (grant #74105 awarded to M.P.B.). This work was also supported by Canadian Institutes of Health Research Postdoctoral Fellowships awarded to JQL and ATK, Fonds de Recherche du Québec Postdoctoral Fellowship awarded to JQL, and the Natural Sciences and Engineering Research Council of Canada Banting Fellowship awarded to ATK.

## AUTHOR CONTRIBUTIONS

Conceptualization: JQL, ATK, MPB

Methodology: JQL, ATK

Investigation: JQL, ATK, EC

Visualization: JQL, ATK

Funding acquisition: JQL, ATK, MPB

Project administration: JQL, ATK, MPB

Supervision: MPB

Writing – original draft: JQL, ATK

Writing – review & editing: JQL, ATK, MPB

## METHODS

### RESOURCE AVAILABILITY

#### Lead contact

Further information and requests should be directed to the lead contacts, J. Quinn Lee and Mark P. Brandon.

#### Materials availability

This study did not generate new unique reagents.

#### Data and code availability

- All data to reproduce the figures will be available upon the date of publication at https://github.com/jquinnlee/weirdgeos.
- All original code will be deposited at Zenodo and will be publicly available at the date of publication.

## EXPERIMENTAL MODEL AND SUBJECT DETAILS

### Animals

Naive male (4) and female (3) mice (C57Bl/6, Charles River) were housed in pairs on a 12-hour light/dark cycle at 22°C and 40% humidity with food and water ad libitum. All experiments were carried out during the light portion of the light/dark cycle, and in accordance with McGill University and Douglas Hospital Research Centre Animal Use and Care Committee (protocol #20157725) and with Canadian Institutes of Health Research guidelines.

## METHOD DETAILS

### Surgical procedures

During all surgeries, mice were anesthetized via inhalation of a combination of oxygen and 4% Isoflurane before being transferred to the stereotaxic frame (David Kopf Instruments), where anesthesia was maintained via inhalation of oxygen and 0.5-2.5% Isoflurane for the duration of the surgery. Body temperature was maintained with a heating pad and eyes were hydrated with gel (Optixcare). Carprofen (10 ml kg^-1^) and saline (0.5 ml) were administered subcutaneously at the beginning of each surgery. Preparation for recordings involved three surgeries per mouse.

First, at the age of six to ten weeks, each mouse was transfected with a 500 nl injection of the calcium reporter GCaMP6f via the viral construct AAV9.syn.GCaMP6f.WPRE.SV40. The original titre of the AAV9.syn.GCaMP6f.WPRE.SV40 construct, sourced from University of Pennsylvania Vector Core, was 3.26e14 GC-ml and was diluted in sterile PBS at a 1:30 ratio before surgical microinjection.

Between five to six weeks post-injection, a 1.8mm diameter gradient refractive index (GRIN) lens (Edmund Optics) was implanted above dorsal CA1 (Referenced to bregma: +/-1.45 mm ML, –1.95 mm AP; –1.37 mm DV from skull surface). Implantation required aspiration of intervening cortical tissue. In addition to the GRIN lens, three stainless steel screws were threaded into the frontal, occipital and parietal bone (above the contralateral hippocampus) to stabilize the implant. Dental cement (C&B Metabond) was applied to secure the GRIN lens and anchor screws to the skull. A silicone adhesive (Kwik-Sil, World Precision Instruments) was applied to protect the top surface of the GRIN lens until the next surgery.

Two weeks after lens implantation, an aluminum baseplate was affixed via dental cement (C&B Metabond) to the skull of the mouse, which would later secure the miniaturized fluorescent endoscope (miniscope) in place during recording. The miniscope/baseplate was mounted to a stereotaxic arm for lowering above the implanted GRIN lens until the field of view contained visible cell segments and dental cement was applied to affix the baseplate to the skull. A polyoxymethylene cap with a metal nut weighing ∼3 g was affixed to the baseplate when the mice were not being recorded, to protect the baseplate and lens, as well as to simulate the weight of the miniscope.

After surgery, animals were continuously monitored until they recovered. For the initial three days after surgery mice were provided with a soft diet supplemented with Carprofen for pain management (MediGel CPF, ∼5mg kg^-1^ each day). Familiarization with the open square environment began at least 1 week following the baseplate surgery.

### Data acquisition

*In vivo* calcium videos were recorded with a UCLA miniscope (v3; miniscope.org) containing a monochrome CMOS imaging sensor (MT9V032C12STM, ON Semiconductor) connected to a custom data acquisition (DAQ) box (miniscope.org) with a lightweight, flexible coaxial cable. The DAQ was connected to a PC with a USB 3.0 SuperSpeed cable and controlled with Miniscope custom acquisition software (miniscope.org). The outgoing excitation LED was set to between 2-8% (∼0.05-0.2 mW), depending on the mouse to maximize signal quality with the minimum possible excitation light to mitigate the risk of photobleaching. Gain was adjusted to match the dynamic range of the recorded video to the fluctuations of the calcium signal for each recording to avoid saturation. Behavioral video data were recorded by a webcam mounted above the environment. The DAQ simultaneously acquired behavioral and cellular imaging streams at 30 Hz as uncompressed AVI files and all recorded frames were timestamped for post-hoc alignment.

All recording environments were constructed of white Lego base plates (Lego, Inc) and white painted plywood walls. The full square environment was 75 cm x 75 cm. The external walls were 60 cm tall, while inserted partition walls were 25 cm tall. A blue Lego baseplate was affixed at the top-center of the Eastern wall to serve as an orienting visual feature. During recording, the environment was dimly lit. All sessions were 40 min, and one session was recorded per day to avoid photobleaching. The order of the environmental configurations was randomized for each mouse, but repeated across sequences such that repeated configurations were always equidistant in time.

### Data preprocessing

Calcium imaging data were preprocessed prior to analyses via a pipeline of open-source MATLAB (MathWorks; version R2015a) functions to correct for motion artifacts^42^, segment cells and extract transients^43,44^. To extract the rising phase of transients from each filtered calcium trace, we proceeded as follows. First, we computed the derivative of the calcium signal, smoothed with a gaussian kernel with a standard deviation of 5 frames. Next, because calcium transients around the baseline can only be positive, we estimated the variance in the derivative of the smoothed calcium signal on the basis of noise via a half-normal distribution such that:

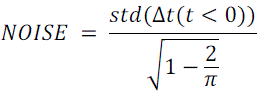

Where Δ*t* is the smoothed time-derivative of the median-subtracted calcium trace *t*. We then z-scored Δ*t* on the basis of this noise distribution. The final binarized rising-phase vector was then set to 1 whenever this z-scored Δ*t* vector exceeded 2.5, and 0 otherwise. This binary vector was treated as the *firing rate* in all further analyses. The motion-corrected calcium imaging data were manually inspected to ensure that motion correction was effective and did not introduce additional artifacts. Following this preprocessing pipeline, the spatial footprints of all cells were manually verified to remove lens artifacts. Position data were inferred from the onboard miniscope red LED offline following recording using a custom written MATLAB (MathWorks) script and were manually corrected if needed. Cells were tracked across sessions on the basis of prominent brain surface landmarks, their spatial footprints, and/or centroids.

### Data analysis

All analyses were conducted using the binary vector of the rising phases of transients, treating this vector as if it were the firing rate of the cell (henceforth *firing rate*). Similar results are observed when the likelihood of spiking was inferred via a second-order autoregressive deconvolution instead of transient rising-phase extraction. All statistical comparisons were performed with Scipy and Statsmodels libraries in Python 3.8.

### Place cell identification

To detect place cells, we generated rate maps for each session half (first and second 20 min) and compared the resulting Pearson correlation to the maps generated from 1000 circular shuffles or the corresponding position data. Place cells were identified as those whose split-half rate map correlation exceeded the 99^th^ percentile of the corresponding shuffled distribution for each cell.

### Bayesian decoding of animal position

To decode animals’ position from calcium traces within each session, we performed a 5-fold split of spatially binned position and trace data and transformed the binned positions to a one-hot vector. Using the Gaussian Naïve Bayes method in the scikit-learn Python library, we predicted position on withheld data from maximum likelihood estimation data following:

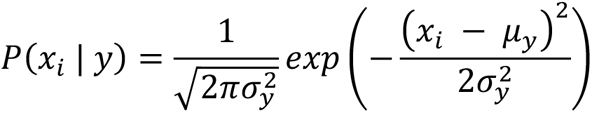

where *x_i_* corresponds to the predicted position at *i* and *y* to the respective calcium traces. We assumed a flat prior (equal likelihood at all positions) and the scikit-learn default σ value of 1e-9. Decoding error was then estimated as the Euclidean distance between the predicted and actual spatial bin position of the animal on withheld data.

### Vector field map generation

To visualize the directional flow of spatial mapping across environments, we calculated the rate map correlation across all cells (i.e., the population vector) for each spatial bin in all geometries relative to the square environment, treating each spatial bin in the square as a “seed” location. We then measured the vector between each spatial bin in the square environment and the maximally correlated bin in each geometry, averaged across animals and sequences. To visualize the spatial displacement of the population vector in each spatial bin, we generated streamplots showing the flow of the population vector fields with matplotlib software (version 3.8.4) colored by the vector magnitude between seed locations in spatial bins of the square environment to the maximally correlated spatial bin in each geometry.

### Representational similarity matrix (RSM) generation

Rate maps were first constructed by spatially binning position data into pixels corresponding to a 5cm x 5cm grid of locations and smoothed with a 5cm standard deviation isometric Gaussian kernel. We then divided rate maps into 9 partitions according to the grid-design of the environment and calculated the pairwise Pearson correlation between all rate maps registered across all pairs of environments and computed the average rate map correlation across all registered cells in each geometry and partition. The resulting RSM was then organized to match the order of geometries and partitions across animals based on the sequence presented to the first subject and in ascending order of partitions from left to right (West to East), and top to bottom (North to South), including each geometry from a respective sequence, starting and ending with the square (Sequence 1: Session 1 – 11; Sequence 2: Session 11 – 21; Sequence 3: Session 21 – 31).

### Non-metric multi-dimensional scaling (nMDS)

Embeddings of the population representation via nMDS were computed as follows: First, we computed the mean rate map correlation across cells for each pairwise comparison of partitions, as described in the main text. Next, we transformed this correlation matrix into a distance matrix by computing one minus this correlation matrix, with the diagonal set to 0 distance. Finally, this distance matrix was reduced to a two-dimensional embedding via nonmetric multidimensional scaling, implemented by the MDS function in the scikit-learn Python library with conservative parameterizations (*eps*=1e-12, *max_iter*=1e9) and minimizing Kruskal’s normalized stress1 cost function:

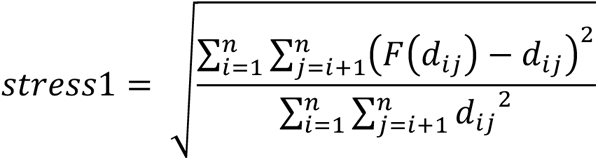

where *d_i,j_* is the distance between *i* and *j* in the measured distance matrix and F(*d_i,j_*) is a nonparametric monotonic function of the measured distances fit via isotonic regression. A score of less than 0.15 is typically considered to indicate a good quality fit.

### Noise ceiling calculation

To estimate the suggested theoretical maximum level of model prediction we calculated the upper and lower bounds of the noise ceiling across animals and sequences. To calculate the lower bound, we excluded one RSM at a time from the averaged RSM calculated across animals and sequences, and then computed the rank-order correlation between the excluded RSM and the remaining RSMs averaged across animals and sequences. To calculate the noise ceiling upper bound, we calculated the average rank-order correlation of each RSM to the average of all RSMs across animals and sequences. This range of values between the upper and lower bound thus indicate the average correlation of an unseen animal with the group representation.

### Modelling of spatial cell types

To model spatial features in the parahippocampal system according to each theoretical view, we utilized a recently developed open-source toolbox^36^ and custom functions in Python 3.8. Parameterizations for each model were based on those defined in previous work^30–35^. In addition, for each simulated cell type we drew firing rates from the modelled spatial receptive fields using animal trajectories from each recorded session recorded at 30 Hz.

### Boundary-vector cells (BVC)

We generated a population of BVCs according to same methods used in prior studies^31–34^. As described previously^34^, the firing rate of the *i*^th^ BVC, with preferred distance *d*_*i*_ and angle φ_*i*_ to a boundary at distance r and direction θ subtending at angle δθ is given by:

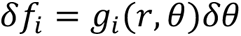

where,

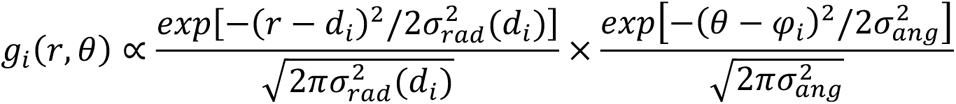

in which the angular tuning width σ*_ang_*, is constant and the radial tuning increases with the preferred distance tuning 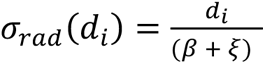 for constants β and ξ. To define the tuning distances of BVCs, we randomly drew values from 0 and 75 cm from a β distribution with parameters α and β between 0.25 and 5.0, wherein α and β both equal to 1.0 generate a random uniform distribution of tuning distances.

### Boundary-vector to place cells (BVC2PC)

According to the boundary-vector to place model^31–33^, we calculated the firing of a model place cell from the input from at least 2 and up to 16 model BVCs. Specifically, we calculated the geometric mean of randomly selected BVC inputs as in Grieves et al.^33^:

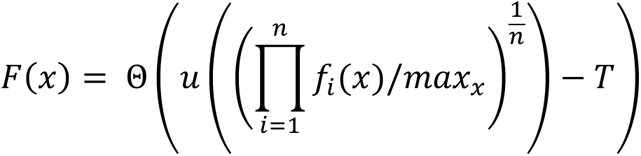

where *T* is the cell’s threshold, defined as 80 percent of the maximum within-session summed inputs, and

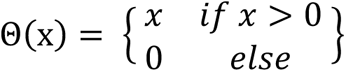

### Boundary-vector to successor features (BVC2SF)

As described previously^34^, we generated BVC2SFs from a population of 200 model BVCs randomly drawn for each animal and learned predictive weights among BVC firing vectors observed through time, according to the temporal difference learning rule:

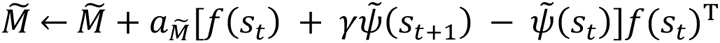

where 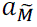 is the learning rate for the successor representation 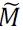 weight vector, and 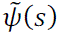 is the expected sum of future BVC population firing rate vectors, discounted exponentially into the future with the parameter γ ∈ [0, 1]. Based on a grid search for approximately optimal parameterizations with a uniform BVC tuning distribution, we set 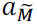 = 0.0167 and γ = 0.999 where *dt* = 1/30. The firing rate of each simulated feature F*_i_* in a location *s* was proportional to the threshold, weighted sum of the BVCs connected to it:

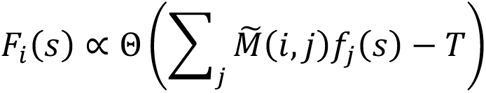

where *T* is the cell’s threshold, and

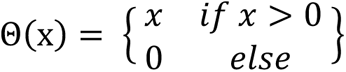

### Grid cells

We modelled the firing of populations of grid cells as the summation of three shifted cosines, as described previously^30^. Briefly, grid cell functions were constructed from the sum of three two-dimensional sinusoidal gratings with 60– and 120-degree angular differences, and firing rates between 0 and 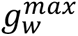, given by:

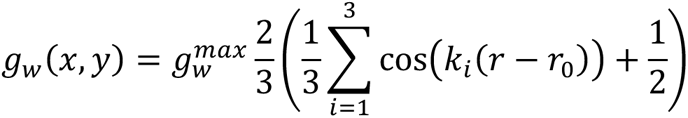

where *r* defines the spatial phase, and *k*_*i*_ defines the wave vectors at *i*.

### Grid to place cell (GC2PC model)

As described previously, we simulated individual place cells from the input of 50 randomly generated grid cells with a single phase, and logarithmic grid-spacing between 28 cm to 73 cm and uniformly random grid orientations. Grid inputs to each place cell were then defined according to the weight coefficient:

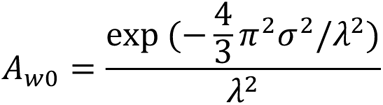

where λ is the grid spacing of the cell’s respective inputs, and the constant σ = 12 cm, as in Solstad et al.^30^. While the primary results from the prior study did not explicitly define the extent inhibition is to be applied to the grid cell inputs, we modelled local inhibition with the summed a threshold *T*, defined as 80% of the maximum within-session firing rates of grid inputs. Inhibition was then applied to the same GC2PC prediction (GC2PC-th), and place cell firing was estimated as:

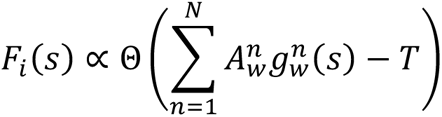

where,

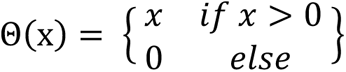

For model features without inhibition, the same procedure was applied without the subtraction of the threshold *T*.

### Boundary-tethered grid to place cells (bt-GC2PC)

To simulate a grid to place cell model with dynamic boundary-tethering^15^, we adopted a parameter-free approach to boundary tethering of the same GC2PC features described above, as in Solstad et al.^30^. To this end, we first generated boundary-tethered receptive fields for the North, South, East, and West boundaries by translating the GC2PC fields generated in the square environment by their relative distance to the walls of each geometry. Following contact with the respective wall (animal within 6 cm of boundary), we sampled from the respective boundary-tethered receptive field. bt-GC2PC features were calculated as the average of the sampled boundary-tethered maps, and inhibition was applied identically as described above (bt-GC2PC-th).

### Naïve place cells (PC)

We simulated naïve place cells (PC) as two-dimensional gaussian receptive fields with a maximum firing rate of 1 Hz and a field width of 20 cm, randomly and uniformly distributed throughout the environment.

### Place to successor features (PC2SF)

We also generated successor features from the naïve place cell (PC) basis set described above with the same learning parameters as the BVC2SF model.

### Supplementary Figures

**Figure S1.**
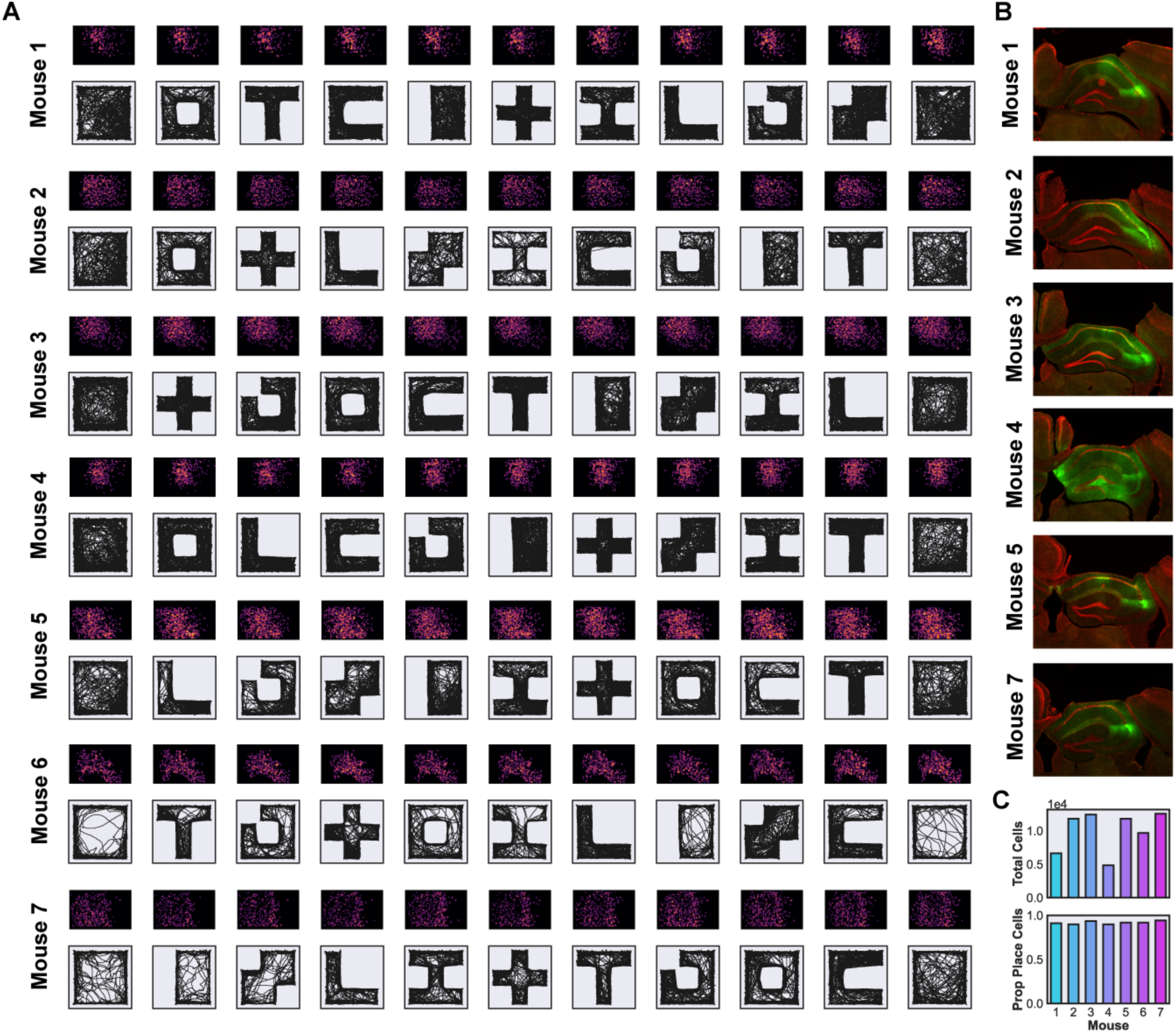
**Recording large neuronal populations with miniscope calcium imaging in freely behaving mice** (**A**) Examples of spatial footprint (SFP) max-projections across a single geometric sequence in each animal following source extraction and across session image alignment (top), along with the respective animal’s estimated position using Deeplabcut (head center; bottom) (*46*). (**B**) Histological confirmation of GRIN lens implantation above dorsal CA1 (DAPI in red, GCaMP6f in green). Tissue damage following post-mortem lens extraction prevented histological confirmation of CA1 implant location in Mouse 6, but subsequent analyses confirmed this mouse was similar to the remaining subjects in the group (see below). (**C**) Top panel shows the total number of cells detected across all sessions in each animal (non-unique cells prior to across-session registration), and the bottom panel shows that the detected proportion of place cells did not differ across mice, confirming similar quality of spatial coding across subjects in CA1.

**Figure S2.**
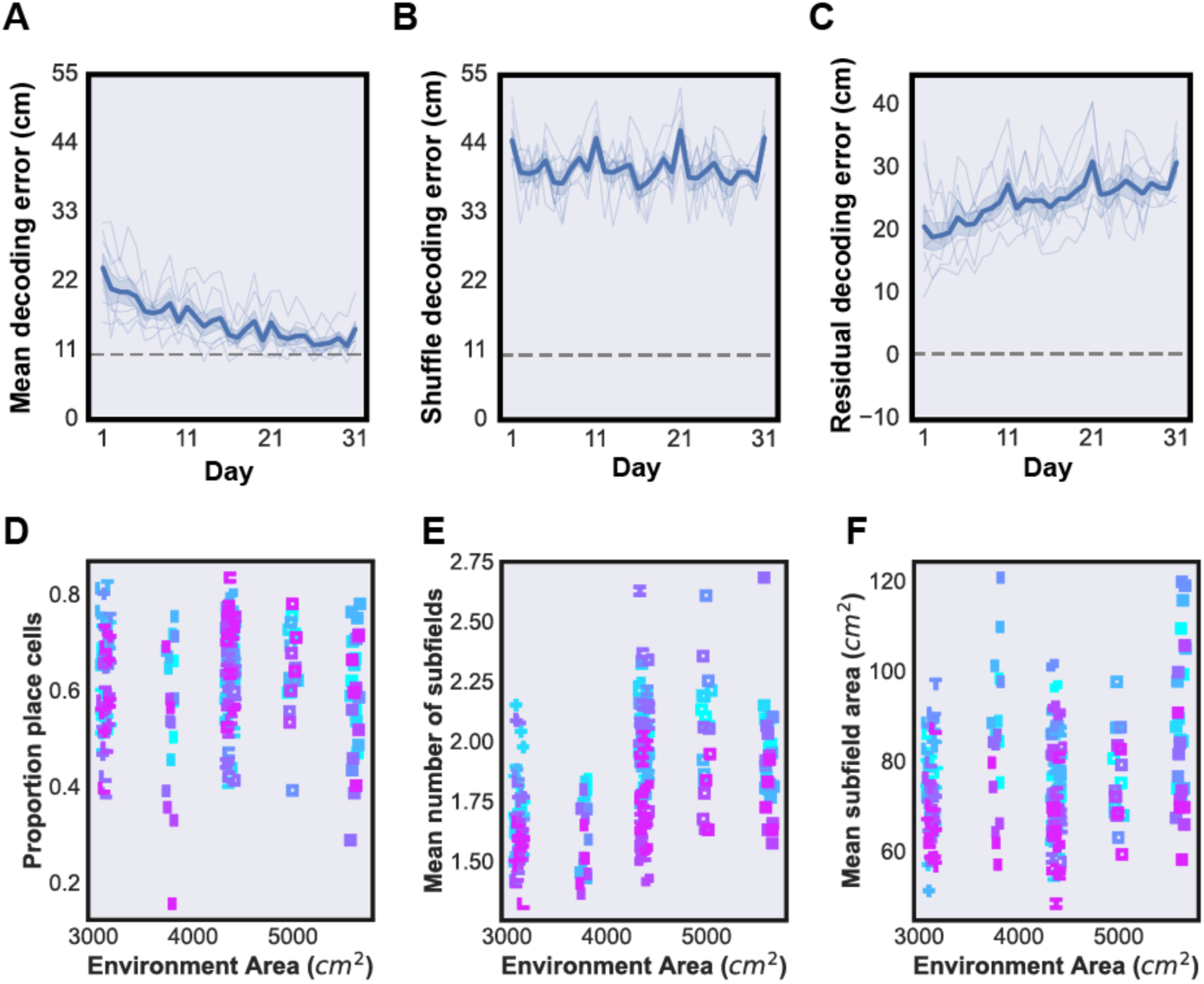
Precise spatial coding of dynamic environments in CA1 across protracted experience (**A**) To confirm previously observed decreases in decoding error across days, we down-sampled the number of cells to match the minimum in each animal registered across all sessions, and predicted each animal’s position with a Bayesian decoder (STAR Methods). Complementing our results with the inclusion of all cells, we found a similar decrease in decoding error across sessions (ANOVA: *P*(Day)<0.0001, *F*(Day)=4.2184). The gray line indicates anticipated theoretical maximum level of decoding accuracy (10 cm) based on previous computational work (*21*). (**B**) Circular shuffling procedures illustrate chance levels of decoding accuracy across all sessions. (**C**) To calculate the residual decoding error, we subtracted the mean decoding error from circularly shuffled data for each animal, and found the residual decoding error improved across sessions, and was significantly above chance levels (ANOVA: *P*(Day)=0.0016, *F*(Day)=2.1055, One-sample t-test against chance level: *P*<0.0001, *t*=59.5178). (**D-F**) Following detection of place cells across all days, we compared the proportion of place cells, mean number of detected subfields (STAR Methods), and subfield area to varied environment area across all geometries and animals. Each datum shows the recorded geometry in the dot shape, and the animal in dot color. We found no effect of environment area on the proportion of detected place cells (Pearson correlation: *P*=0.4289, *R*=0.0640), but did observe a significant effect on the number of detected subfields (*P*<0.0001, *R*=0.4538), and subfield area for detected place cells (*P*<0.0001, *R*=0.3358).

**Figure S3.**
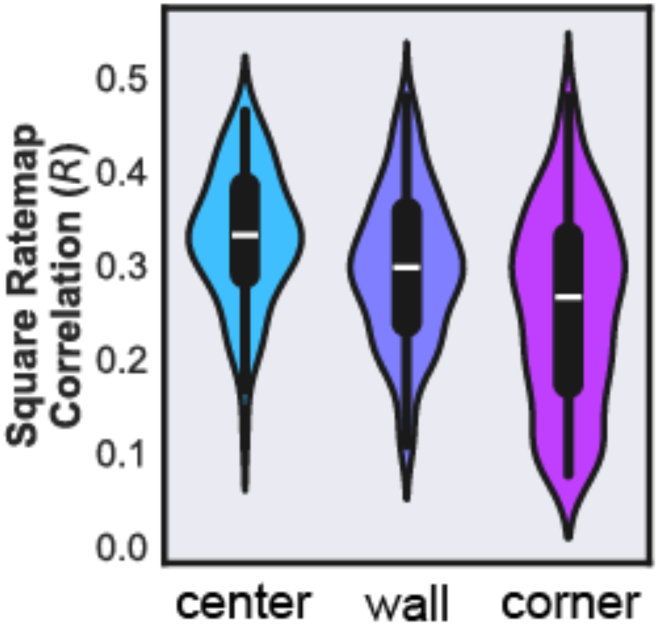
Effect of partition occlusions by location on remapping To determine the relative impact of occluding different regions of the environment on remapping in CA1, we calculated the similarity of rate maps in each geometry to the square environment when environmental deformations included omission of the center, wall, or corner partitions. A one-way ANOVA revealed a significant effect of occlusion location on rate map similarity, wherein occlusion of corner partitions induced the most significant amount of remapping, followed by wall and center partition occlusion (*P*(Occlusion)<0.0001, *F*(Occlusion)= 24.5685).

**Figure S4.**
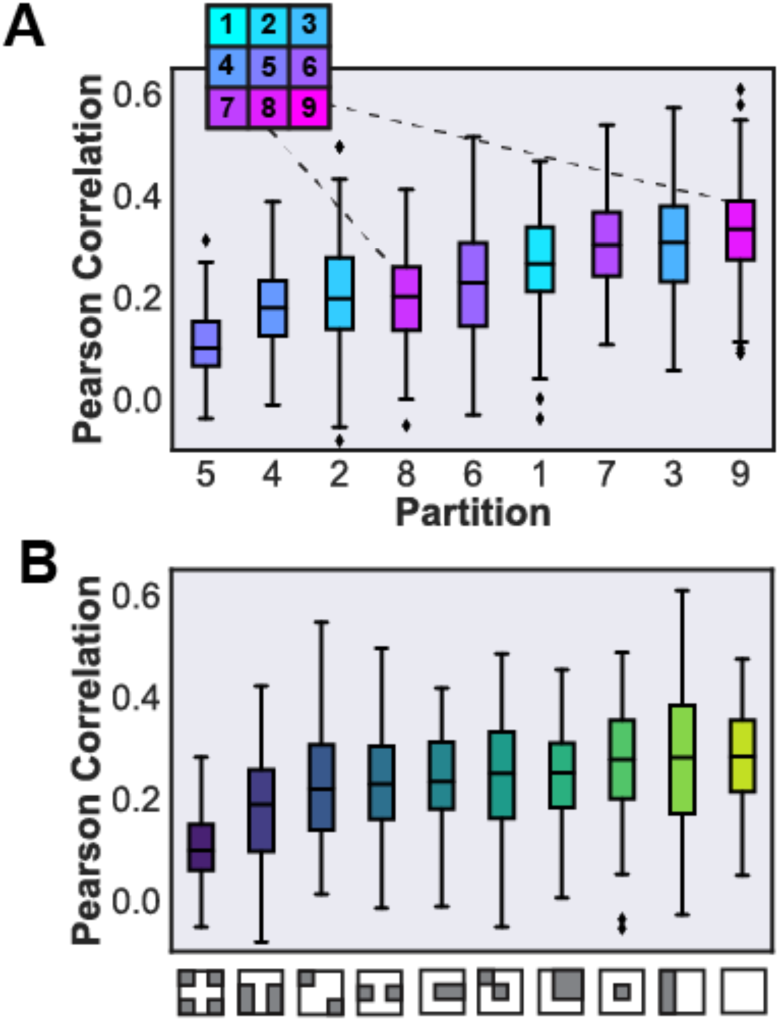
Rate map correlation across partitions and geometries (**A**) Similarity of individual partitions of all deformed geometries to the matched partition in the square (ANOVA: *P*(Partition)<0.0001, *F*(Partition)=102.2067). (**B**) All partitions in deformed geometries relative to the same partition in the square for each environment (ANOVA: *P*(Geometry)<0.0001, *F*(Geometry)=29.3338; *P*(Geometry X Partition)<0.0001, *F*(Geometry X Partition)=2.8847).

**Figure S5.**
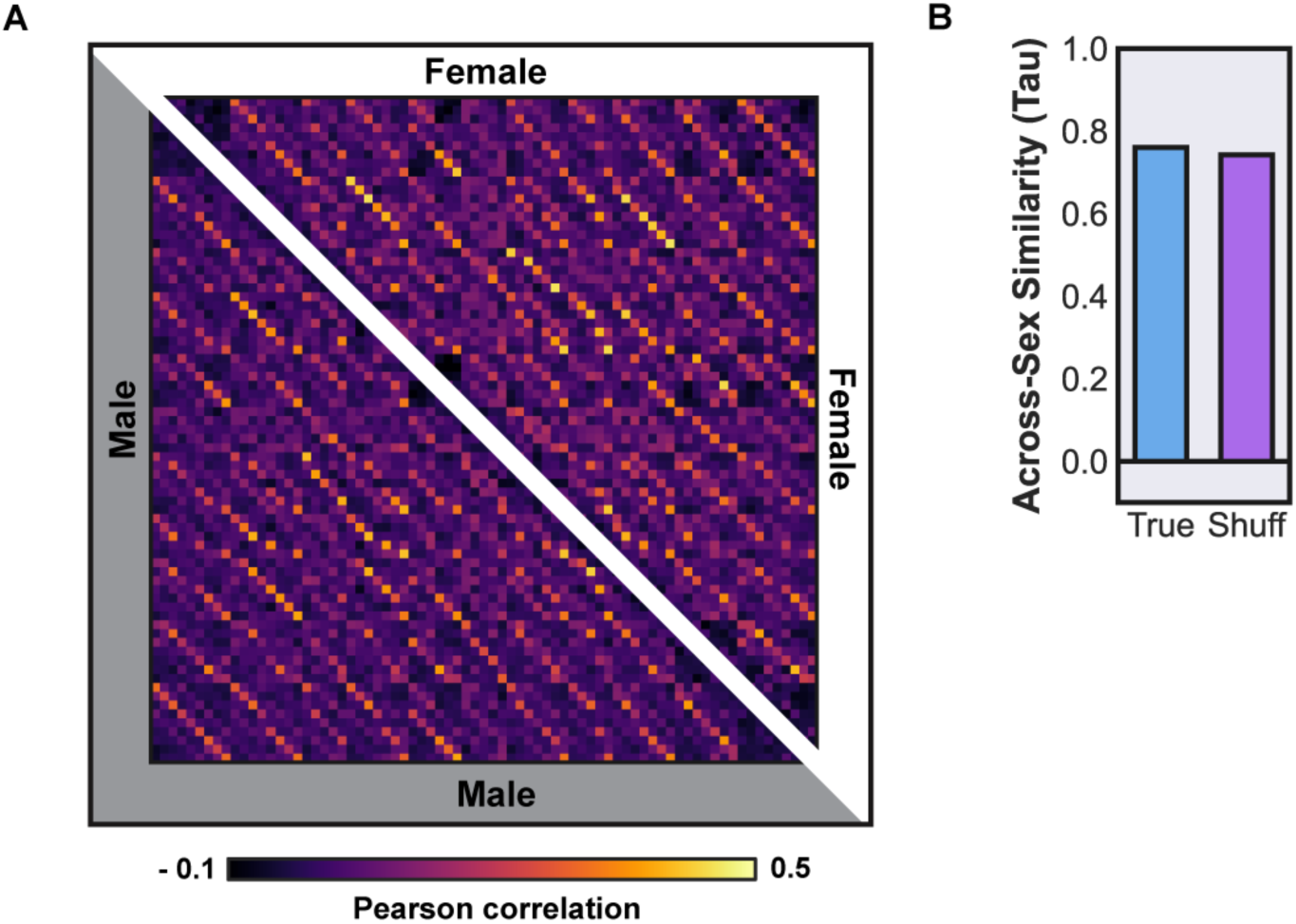
Sex does not determine the representation of geometrically dynamic environments in CA1 (**A**) The RSM shows the averaged representation of male and female mice across all sequences in CA1. (**B**) To determine whether sex affects task representation, we calculated the rank-order correlation (Kendall’s Tau) of the aggregate male and female RSM, compared to the average of 100 shuffles of sex ID. We found no difference between male and female RSM correlation compared to the shuffled controls of sex (One-sample t-test: *P*=0.9999, *t*=-5.6237).

**Figure S6.**
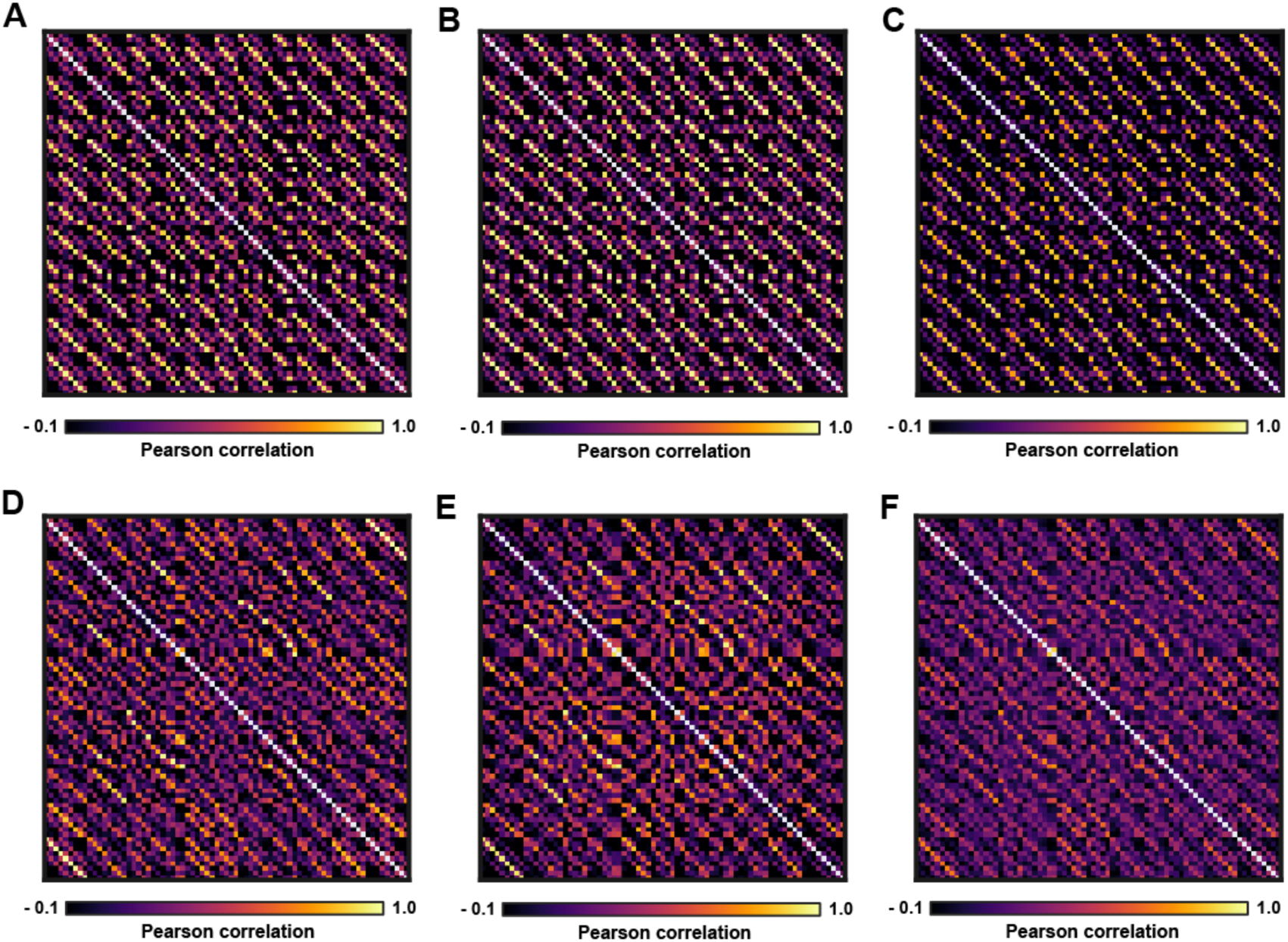
RSMs for CA1 data and theoretical model predictions (**A**) RSM for the PC model. (**B**) RSM for the GC2PC model. (**C**) RSM for the PC2SF model. (**D**) RSM for the bt-GC2PC model. (**E**) RSM for the BVC2PC model. (**F**) RSM for the BVC2SF model.

**Figure S7.**
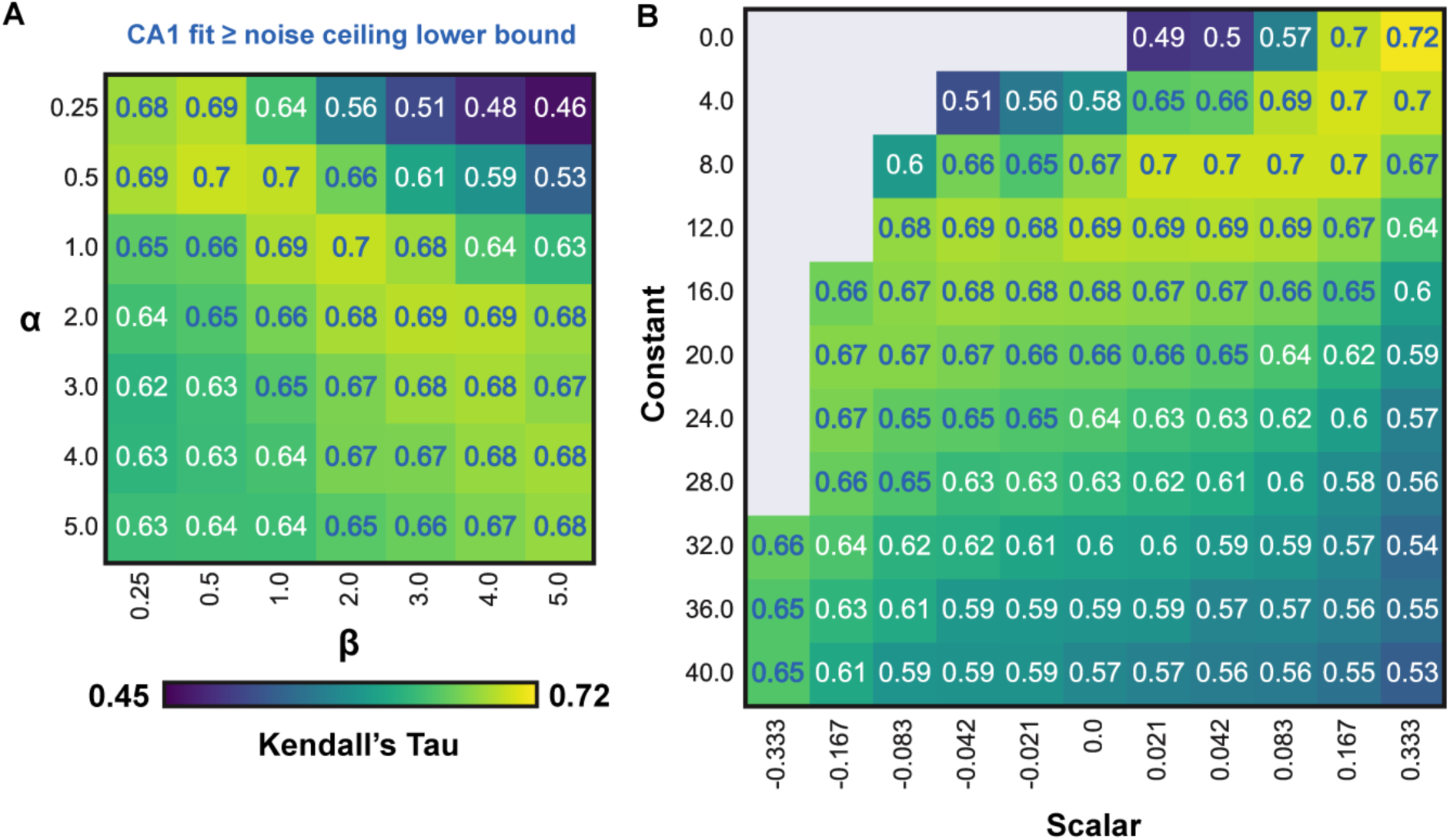
BVC2PC model grid search with prediction accuracy for CA1 representation (**A**) Resulting model fit to CA1 (Kendall’s Tau) for alpha and beta distribution parameters for BVC preferred tuning distances (*N* = 49). White labels indicate model parameterizations that fall below the noise ceiling lower bound, while blue labels indicate model parameters that result in predicted representations above the noise ceiling lower bound. (**B**) Model fit to CA1 (Kendall’s Tau) for BVC distance constant and scalar parameterizations, wherein blue labels indicate model parameterizations above noise ceiling lower bound (*N* = 104).

**Figure S8.**
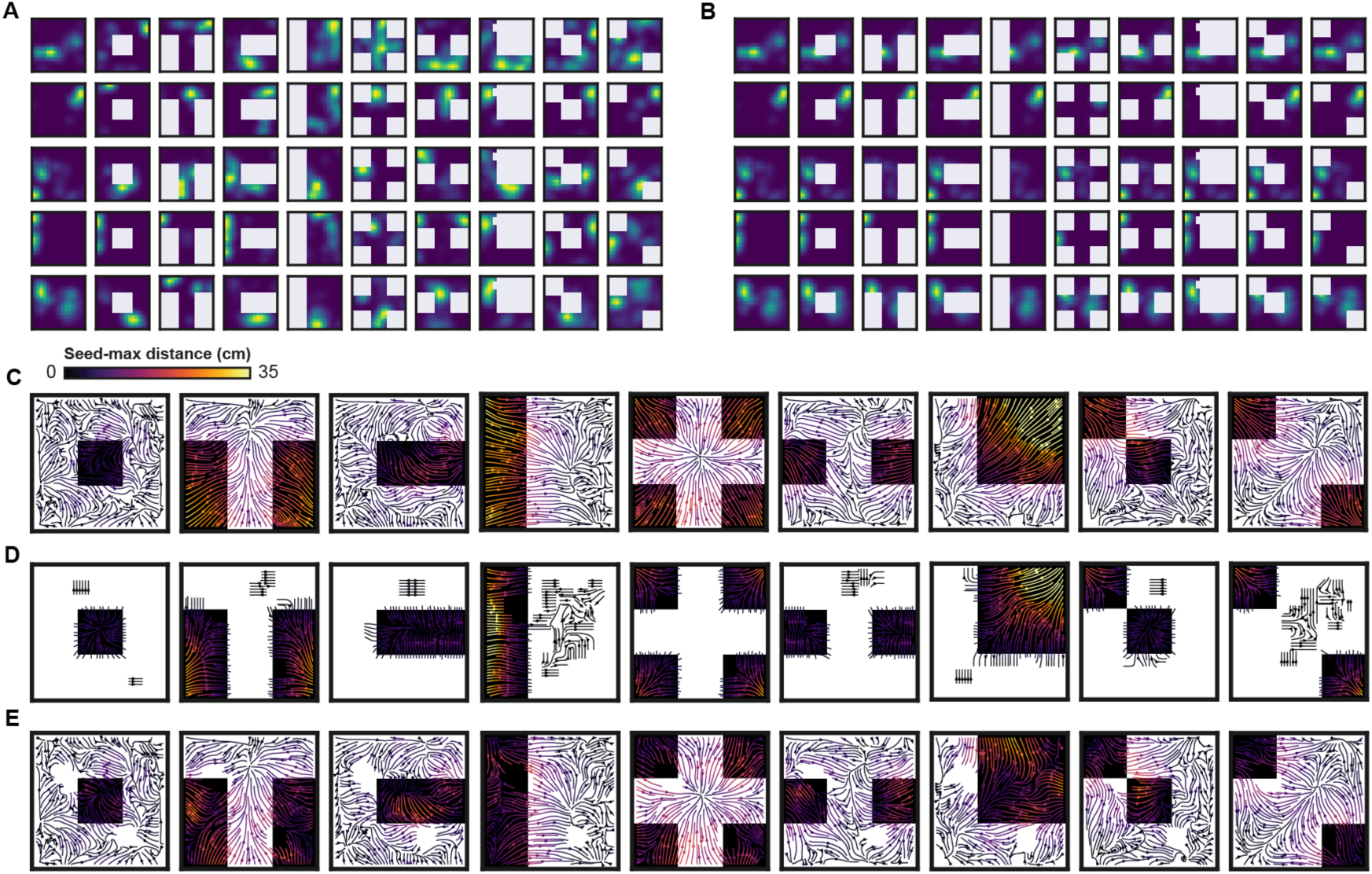
Geometric remapping with simulated spatial stability and environment partitioning To determine whether similar vector fields would be observed due to partitioning of environments with allocentrically static spatial mapping, we simulated stable maps using rate maps from cells active in the first square environment, and masked regions sampled by the animal in each session. We then calculated vector fields from simulated maps, which revealed anticipated movement near boundaries, but lack of vector fields in sampled partitions. (**A**) Example rate maps from CA1 across geometries. (**B**) The same rate maps shown in (B) following the stability simulation with masking. (**C**) Vector field stream plots from actual rate maps for each geometry compared to the square. (**D**) Vector field stream plots from stable simulation masked rate maps, such as those shown in (B). (**E**) To confirm the effect of geometric remapping, we calculated the residual vector fields by subtracting those from stable field simulation and partitioning them from vector fields calculated with actual data. With this approach, we continued to observe a strong effect of geometry on the residual vector fields, confirming the effect of geometric remapping observed in CA1. The figure shows vector field stream plots of the residuals.

**Figure S9.**
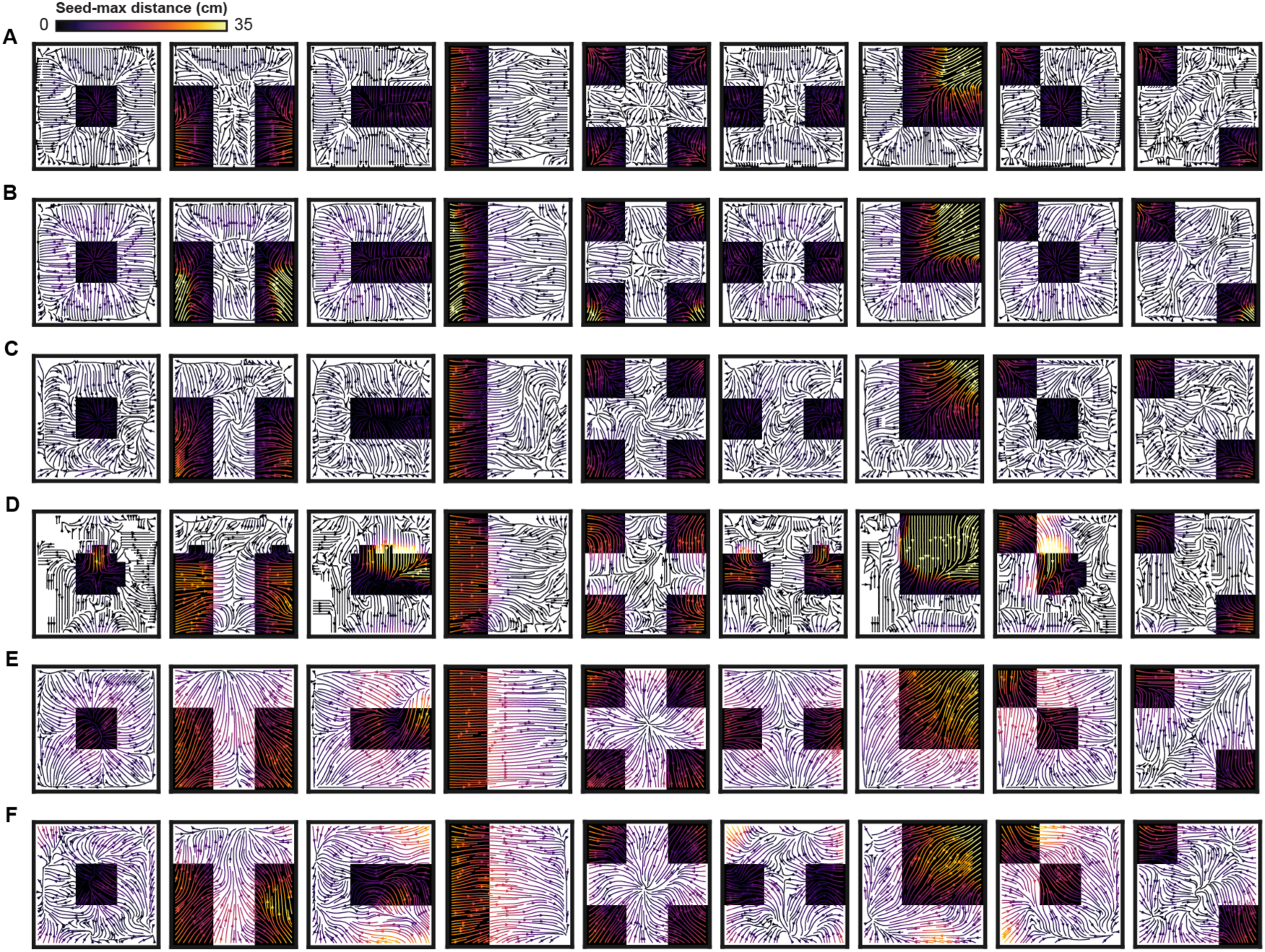
Model-based vector field stream plots (A) PC model. (B) GC2PC model. (C) PC2SF model. (D) bt-GC2PC model. (E) BVC2PC model. (F) BVC2SF model.

